# GLP1R agonists and vorasidenib converge on peroxynitrite-mediated cytotoxicity in mutant IDH glioma

**DOI:** 10.64898/2026.10.18.759957

**Authors:** Céline Taglang, Suresh Udutha, Timothy Phoenix, Justin D. Lathia, John de Groot, Pavithra Viswanath

**Affiliations:** Department of Radiology and Biomedical Imaging, University of California San Francisco, San Francisco, CA, USA; Division of Pharmaceutical Sciences, University of Cincinnati, College of Pharmacy, Cincinnati, OH, USA; Department of Cancer Sciences, Cleveland Clinic Research, Cleveland, OH, USA; Department of Molecular Medicine, Cleveland Clinic Lerner College of Medicine, Case Western Reserve University, Cleveland, OH, USA; Case Comprehensive Cancer Center, Cleveland, OH, USA; Rose Ella Burkhardt Brain Tumor & Neuro-Oncology Center, Cleveland, OH, USA; Department of Biological, Geological, and Environmental Sciences, Cleveland State University, Cleveland, OH, USA; Department of Clinical Medicine, Faculty of Health and Medical Sciences, University of Copenhagen, Denmark; Department of Neurology and Neurological Surgery, University of California San Francisco, San Francisco, CA, USA

## Abstract

Mutations in isocitrate dehydrogenase 1 or 2 (mIDH) generate the oncometabolite D-2-hydroxyglutarate that drives tumorigenesis in a distinct molecular class of gliomas. The mIDH inhibitor vorasidenib slows tumor growth, but disease progression remains a challenge, identifying a need for strategies that deepen and prolong therapeutic response. Here, we show that glucagon-like peptide-1R (GLP1R) agonists synergize with vorasidenib in patient-derived and murine models of mIDH glioma. Mechanistically, GLP1R agonists activate cAMP-protein kinase A signaling and induce nitric oxide synthase 1-dependent nitric oxide production in mIDH glioma cells. Nitric oxide inhibits aconitase and ribonucleotide reductase via S-nitrosylation, thereby disrupting the tricarboxylic acid cycle, depleting deoxyribonucleotides, blocking DNA synthesis, and exerting a cytostatic inhibition of tumor cell proliferation. Concomitantly, vorasidenib relieves D-2HG-driven suppression of pyruvate dehydrogenase, thereby increasing mitochondrial superoxide production and suppressing proliferation without cytotoxicity. When combined, GLP1R agonist-induced nitric oxide and vorasidenib-induced superoxide converge to generate peroxynitrite, which damages lipids and DNA, and triggers apoptosis in mIDH glioma cells. Combining a GLP1R agonist with vorasidenib recapitulates these metabolic and pharmacodynamic effects in mice bearing intracranial mIDH tumors, induces tumor regression, and significantly prolongs survival relative to each monotherapy in vivo. Collectively, our studies demonstrate that GLP1R agonism and mIDH inhibition induce distinct alterations in mitochondrial redox metabolism that converge to produce synergistic cytotoxicity and provide a mechanistic basis for clinical evaluation of this therapeutic strategy in patients with mIDH glioma.

## INTRODUCTION

Diffuse gliomas are the most common form of malignant primary brain cancer in adults^1^. The presence of a mutation in isocitrate dehydrogenase 1 or 2 (mIDH) is the first diagnostic criterion for diffuse gliomas^2^. Tumors that are isocitrate dehydrogenase wild type are classified as glioblastoma (GBM, grade 4)^2,3^. mIDH gliomas are further divided into oligodendrogliomas (grades 2-3) or astrocytomas (grades 2-4) based on additional genetic alterations^2,3^. Although mIDH gliomas generally grow more slowly and confer longer survival than GBMs, they remain incurable and patients with mIDH gliomas experience profound, life-altering neurological, cognitive, and functional deficits^3,4^.

The mIDH enzyme converts the normal cellular metabolite α-ketoglutarate into the oncometabolite D-2-hydroxyglutarate (D-2HG), which accumulates to millimolar concentrations in these tumors^5^. D-2HG competitively inhibits the activity of >70 α-ketoglutarate-dependent enzymes, including histone and DNA demethylases, leading to widespread epigenetic alterations that block differentiation and drive tumorigenesis^6–8^.

The development of the mIDH inhibitor vorasidenib (VOR) represents a significant milestone in the management of patients with mIDH glioma^9–12^. VOR suppresses D-2HG production and partially restores histone and DNA demethylation in patients with mIDH gliomas^9^. The phase 3 INDIGO trial showed that VOR significantly extends progression-free survival and increases the time to next intervention in patients with grade 2 mIDH gliomas who never received any therapy other than surgical tumor resection^10,11^. These results led to the approval of VOR as a first-line therapy for patients with grade 2 mIDH gliomas and as a potential therapeutic option for patients with grade 3 tumors^12^. While undoubtedly significant, patients eventually experience tumor progression on VOR, highlighting a need for strategies that can deepen and prolong the therapeutic benefit^10,11,13,14^.

D-2HG broadly rewires the metabolome of mIDH glioma cells. We and others have demonstrated that D-2HG alters glucose, glutamine, nucleotide, redox, and phospholipid metabolism in mIDH glioma cells^15–19^. Specifically, we demonstrated that D-2HG inhibits the activity of pyruvate dehydrogenase (PDH), the rate-limiting enzyme in the oxidation of glucose-derived pyruvate via the tricarboxylic acid (TCA) cycle^16^. We also showed that pharmacological activation of PDH was detrimental to the viability of mIDH glioma cells^16^. However, whether VOR reverses D-2HG-mediated inhibition of PDH, and whether this represents a targetable metabolic vulnerability in the context of VOR treatment, remains unknown.

Glucagon-like peptide-1 receptor (GLP1R) agonists are a clinically approved class of drugs that mimic the endogenous gut peptide GLP-1 and have shown remarkable efficacy in the treatment of type 2 diabetes and obesity^20^. The GLP1R is expressed at low levels across multiple cell types and signaling via the GLP1R activates the Gs-adenylyl cyclase-cAMP-protein kinase A (PKA) pathway, leading to modulation of ion channels, intracellular Ca²⁺ signaling, kinase activity, and transcriptional programs^21,22^. These signaling events govern processes such as hormone and neurotransmitter secretion, neuronal excitability, metabolism, survival, and plasticity in a cell-type-specific manner^20–22^. Notably, several of these pathways regulate cellular proliferation and have been implicated in tumor growth. Many tumors, including brain tumors, express GLP1R, and higher GLP1R expression has been associated with improved survival in breast, esophageal, kidney, and thyroid cancers^23^. Retrospective studies suggest that GLP1R agonist use may be associated with reduced incidence and improved survival of patients with obesity-associated cancers^24,25^. The ability of GLP1R agonists to modulate signaling and metabolic pathways linked to tumor proliferation, coupled with these clinical observations, provides a rationale for investigating their activity in mIDH glioma. However, whether mIDH glioma cells express functional GLP1R and whether GLP1R agonists directly alter their growth or therapeutic response is unknown.

The goal of this study was to examine the effects of GLP1R agonists on the metabolism, proliferation, and VOR response of mIDH gliomas. Using clinically relevant patient-derived and murine models^26,27^, we established that mIDH glioma cells express surface-accessible GLP1R. GLP1R agonists, including liraglutide^21^ and NLY01^28^, activated cAMP-PKA signaling and induced mitochondrial nitric oxide (NO) production in a nitric oxide synthase 1 (NOS1)-dependent manner in mIDH glioma cells. NO, in turn, induced S-nitrosylation of TCA cycle and nucleotide biosynthesis enzymes, leading to nucleotide depletion, a block in DNA synthesis, and cytostatic inhibition of tumor proliferation. Simultaneously, VOR activated PDH and caused mitochondrial superoxide generation, also leading to a cytostatic block in tumor proliferation. Importantly, combining a GLP1R agonist with VOR led to the formation of peroxynitrite, which damaged DNA and lipids, triggered apoptotic cell death, and significantly prolonged survival *in vivo*. Our studies, therefore, lay the mechanistic foundation for clinical translation of the combination of GLP1R agonists with VOR in patients with mIDH gliomas.

## RESULTS

### mIDH gliomas express GLP1R on the cell surface

To begin with, we set out to determine whether functional GLP1R expression could be detected in mIDH gliomas. Given the low abundance of GLP1R mRNA in many cell types and the questionable specificity of detecting GLP1R mRNA or protein^29,30^, we quantified surface-accessible GLP1R using live-cell labeling with LUXendin645^31^, a fluorescent GLP1R antagonist, followed by flow cytometry (Fig. 1A). First, we demonstrated that LUXendin645 robustly labeled HEK293 cells expressing GLP1R but gave no signal above the unstained background in isogenic GLP1R-knockout HEK293 cells (Fig. 1A-1B), establishing that the probe provides a specific readout of GLP1R. LUXendin645 labeling was detectable across multiple patient-derived (oligodendroglioma: TS603, SF10417; astrocytoma BT142) and murine (133-A6, 130-A7, and SB28mut) mIDH glioma models (Fig. 1B). Prior incubation with the GLP1R antagonist exendin-(9-39)^32,33^ (EXE) abolished the LUXendin645 signal (Fig. 1B), further confirming the specificity of our readout. Furthermore, LUXendin645 staining did not exceed the unstained background in normal human astrocytes (NHA) or normal human oligodendrocytes (NHO; Fig. 1A-1B).

**Figure 1.**
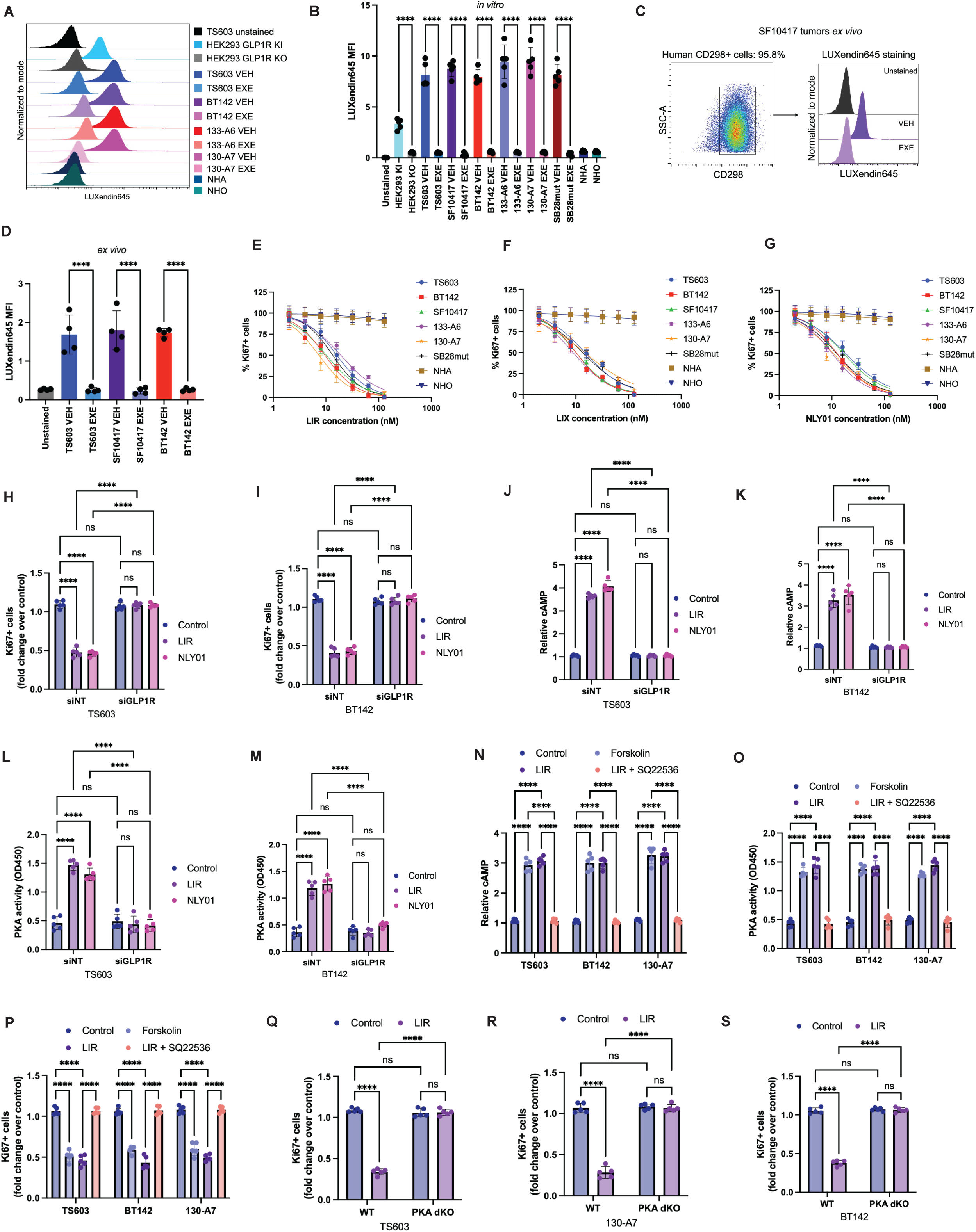
mIDH glioma cells express surface-accessible GLP1R that signals through cAMP-PKA and restrains proliferation. **(A)** Representative live-cell LUXendin645 labeling measured by flow cytometry. Histograms, top to bottom: TS603 unstained; HEK293 GLP1R knock-in (KI) and isogenic GLP1R knockout (KO); TS603, BT142, 133-A6 and 130-A7, each with VEH or exendin-(9-39; EXE) preincubation; NHA; NHO. **(B)** LUXendin645 Median Fluorescence Intensity (MFI) in cells: unstained control, HEK293 GLP1R KI and KO, six mIDH glioma models (TS603, SF10417, BT142, 133-A6, 130-A7, SB28mut) each with VEH or EXE, NHA and NHO. **(C)** Representative live-cell LUXendin645 labeling in intracranial SF10417 tumors analyzed ex vivo. Left, identification of human CD298+ tumor cells (95.8% of events) after depletion of mouse stromal cells. Right, LUXendin645 labeling of the CD298+ gate: unstained, VEH, and EXE. **(D)** LUXendin645 MFI in CD298+ tumor cells: unstained control, and cells recovered from intracranial TS603, SF10417, and BT142 tumors, each with VEH or EXE. **(E-G)** Concentration-response curves for LIR **(E)**, LIX **(F),** and NLY01 **(G)**: Ki67+ cells as a percentage of total, in six mIDH glioma models, NHA and NHO, after 72 h of treatment. **(H, I)** Ki67+ cells (fold change over control) in TS603 **(H)** and BT142 **(I)** transfected with non-targeting (siNT) or GLP1R-directed (siGLP1R) siRNA and treated with vehicle, LIR, or NLY01. **(J, K)** Relative intracellular cAMP in TS603 **(J)** and BT142 **(K)**, siNT versus siGLP1R, treated with vehicle, LIR, or NLY01. **(L, M)** PKA activity (OD450) in TS603 **(L)** and BT142 **(M)**, siNT versus siGLP1R, treated with vehicle, LIR or NLY01. **(N-P)** Relative cAMP **(N)**, PKA activity **(O),** and Ki67+ cells **(P)** in TS603, BT142, and 130-A7 treated with vehicle, the adenylyl cyclase activator forskolin, LIR, or LIR together with the adenylyl cyclase inhibitor SQ22536. **(Q-S)** Ki67+ cells in wild-type and PKA double-knockout (PKA dKO; PRKACA and PRKACB deleted) TS603 **(Q)**, 130-A7 **(R),** and BT142 **(S)** treated with vehicle or LIR. Flow cytometry histograms are normalized to the mode and plotted on an identical axis range. Data are mean ± SD with individual biological replicates shown (n ≥ 3 per group). ****p<0.0001, ***p<0.001, **p<0.01, *p<0.05; ns, not significant.

We next asked whether surface-accessible GLP1R expression was retained in intracranial tumors. Tumors were resected from mice bearing intracranial patient-derived mIDH glioma xenografts and dissociated, and human tumor cells were identified by CD298 expression after depletion of mouse stromal cells (Fig. 1C and Supplementary Fig. S1A). CD298+ tumor cells exhibited LUXendin645 labeling that was reduced to background levels by competition with EXE (Fig. 1D). These results confirm GLP1R expression on the cell surface in mIDH glioma cells and intracranial tumors.

### GLP1R agonists activate cAMP-PKA signaling and inhibit mIDH glioma proliferation in a cytostatic manner

Next, we questioned whether GLP1R expression was functionally relevant in mIDH gliomas. To this end, we treated mIDH glioma cells with liraglutide (LIR), a long-acting acylated human GLP-1 analog; lixisenatide (LIX), a short-acting exendin-4-based agonist; or NLY01, a PEGylated, long-acting exendin-4-based agonist developed for sustained systemic and central nervous system exposure^21,28,34^. All 3 GLP1R agonists robustly inhibited proliferation of a panel of patient-derived and murine mIDH glioma models, with IC50 values of ∼7 to 20 nM (Fig. 1E-1G). In contrast, they failed to significantly affect the proliferation of NHAs or NHOs at concentrations up to 128 nM (Fig. 1E-1G), consistent with the absence of detectable surface GLP1R in these cells (Fig. 1A-1B). To test the specificity of the anti-proliferative effect, we silenced GLP1R in the TS603 and BT142 models. GLP1R silencing did not alter baseline proliferation but abolished the anti-proliferative effect of both agonists (Fig. 1H-1I). Of note, none of the GLP1R agonists induced cell death (Supplementary Fig. S1B), indicating that GLP1R agonism inhibited proliferation in a cytostatic manner.

We then examined whether GLP1R agonists inhibited mIDH glioma proliferation via the classical cAMP-PKA signaling pathway. First, we established that both LIR and NLY01 induced cAMP production (Fig. 1J-1K) and increased PKA activity (Fig. 1L-1M) in mIDH glioma cells, effects that were abolished by GLP1R silencing. To determine whether cAMP-PKA signaling is necessary for the anti-proliferative effect of GLP1R agonists, we examined whether forskolin, an activator of adenylyl cyclase, mimics GLP1R agonism and conversely whether SQ22536, an adenylyl cyclase inhibitor, rescues this effect^35,36^. We confirmed that forskolin increases intracellular cAMP (Fig. 1N) and PKA activity (Fig. 1O) and inhibits proliferation (Fig. 1P) of mIDH glioma cells. Conversely, combining the adenylyl cyclase inhibitor SQ22536 with LIR blocked the LIR-induced increase in cAMP and PKA activity and restored proliferation in mIDH glioma cells (Fig. 1N-1P). Consistent with these pharmacological findings, deletion of both catalytic subunits of PKA (PRKACA and PRKACB; hereafter referred to as PKA-dKO) reduced PKA activity (Supplementary Fig. S1C-S1E) and blocked the anti-proliferative effect of LIR (Fig. 1Q-1S) in the TS603, BT142, and 130-A7 models. Collectively, these results establish that GLP1R agonists suppress the proliferation of mIDH glioma cells in a cytostatic manner through the cAMP-PKA signaling pathway.

### GLP1R agonists inhibit mitochondrial aconitase (ACO2) and ribonucleotide reductase (RNR) in mIDH glioma cells

To define the molecular mechanism driving the anti-proliferative effect, we examined the effect of GLP1R agonists on mIDH glioma metabolism. Treatment with LIR or NLY01 produced a consistent metabolic signature across six patient-derived and murine mIDH glioma models (Fig. 2A-2B). Specifically, both GLP1R agonists depleted the intracellular pool of deoxyribonucleoside triphosphates (dNTPs), isocitrate, α-ketoglutarate, glutamate, and succinate, while increasing citrate and the ribonucleoside diphosphates (NDPs). Because mIDH uses α-ketoglutarate as its substrate, the fall in α-ketoglutarate was accompanied by a reduction in D-2HG. This characteristic metabolic pattern predicted a block at two enzymes, i.e., aconitase 2 (ACO2) in the TCA cycle and ribonucleotide reductase (RNR) in nucleotide biosynthesis (Fig. 2C).

**Figure 2.**
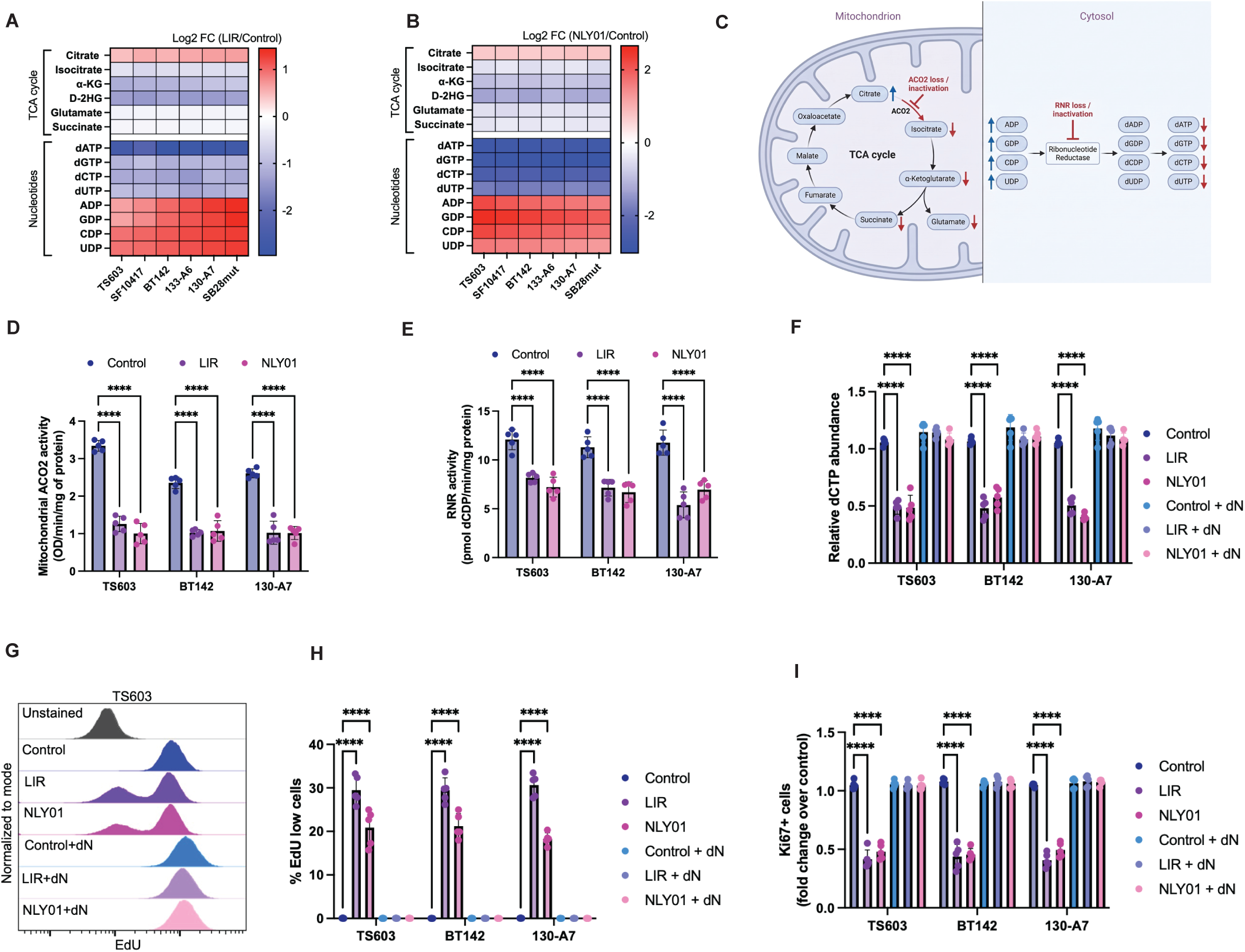
GLP1R agonists inhibit ACO2 and RNR activity and deplete dNTP pools. **(A, B)** Metabolite abundances in six mIDH glioma models treated with LIR **(A)** or NLY01 **(B)** for 72 h, shown as log2 fold change over vehicle. Rows are grouped into TCA cycle intermediates (citrate, isocitrate, alpha-ketoglutarate, D-2HG, glutamate, succinate) and nucleotides (dATP, dGTP, dCTP, dUTP, ADP, GDP, CDP, UDP). Every metabolite differed significantly from vehicle in every model in both panels (84 comparisons per panel, n = 5 per model; the largest p = 0.0014 for LIR and 0.013 for NLY01, and all comparisons remain significant after Holm correction within each panel), so no significance markers are shown. **(C)** Schematic illustration of the metabolic pattern expected following the loss of ACO2 activity in the mitochondrion and the loss of RNR activity in the cytosol. **(D)** Mitochondrial ACO2 activity (OD/min/mg protein) in TS603, BT142, and 130-A7 treated with vehicle, LIR, or NLY01. **(E)** RNR activity (pmol dCDP/min/mg protein) in TS603, BT142, and 130-A7 treated with vehicle, LIR, or NLY01. **(F)** Relative intracellular dCTP abundance in TS603, BT142, and 130-A7 cells treated with vehicle, LIR, or NLY01 with or without deoxyribonucleoside supplementation (dN). **(G)** Representative histograms of EdU incorporation in TS603 cells that were treated with vehicle, LIR, or NLY01 with or without deoxyribonucleoside supplementation (dN). Unstained cells were used as background controls. EdU-low cells as a percentage of total **(H)** and Ki67+ cells (fold change over control; **I**) in TS603, BT142, and 130-A7 cells treated with vehicle, LIR, or NLY01 with or without deoxyribonucleoside supplementation (dN). Flow cytometry histograms are normalized to mode and plotted on an identical axis range. Data are mean ± SD with individual biological replicates shown (n ≥ 3 per group). ****p<0.0001, ***p<0.001, **p<0.01, *p<0.05; ns, not significant.

ACO2 catalyzes the reversible isomerization of citrate to isocitrate through cis-aconitate, and inhibition of ACO2 is expected to result in the accumulation of citrate and the depletion of isocitrate and downstream metabolites, as seen in GLP1R agonist-treated cells. Ribonucleotide reductase (RNR) catalyzes the rate-limiting step in de novo dNTP synthesis by reducing NDPs to deoxyribonucleoside diphosphates (dNDPs), which are subsequently phosphorylated to dNTPs. As such, RNR inhibition is expected to cause the pattern of NDP accumulation and dNTP depletion observed in GLP1R agonist-treated cells. In proliferating cells, active RNR comprises the large RRM1 subunit, which contains the catalytic and allosteric sites, and the small RRM2 subunit, which contains the iron-tyrosyl radical center required to initiate catalysis^37^.

Neither LIR nor NLY01 altered the mRNA or protein abundance of ACO2, RRM1, or RRM2 (Supplementary Fig. S2A-S2D). In contrast, both treatments markedly reduced mitochondrial aconitase and RNR activities across models (Fig. 2D-2E). These results indicated that GLP1R agonism inhibits ACO2 and RNR at the level of enzyme activity rather than expression.

To determine which of the two blocks accounted for the anti-proliferative effect, we bypassed each one independently. dNTPs are essential for DNA synthesis. To assess whether the dNTP depletion induced by GLP1R agonism was functionally relevant, we asked whether GLP1R agonism affected DNA synthesis and whether this could be rescued by supplementation with a mix of deoxyribonucleosides (2′-deoxyadenosine, 2′-deoxyguanosine, 2′-deoxycytidine, thymidine), which bypasses RNR via salvage kinases. First, we confirmed that supplementation with the deoxyribonucleoside mix restored intracellular dCTP abundance in GLP1R agonist-treated mIDH glioma cells (Fig. 2F). Both LIR and NLY01 markedly reduced DNA synthesis, as measured by EdU incorporation, and inhibited proliferation, effects that were rescued by deoxyribonucleoside supplementation (Fig. 2G-2I).

Supplementation with cell-permeable α-ketoglutarate restored the intracellular α-ketoglutarate (Supplementary Fig. S2E) and D-2HG (Supplementary Fig. S2F) pool sizes but failed to restore DNA synthesis (Supplementary Fig. S2G) or proliferation (Fig. S2H) in LIR-treated cells. Each rescue was therefore active on its own arm and inactive on the other. This double dissociation established that the loss of RNR activity and the resulting depletion of dNTPs were sufficient to account for both the block in DNA synthesis and the inhibition of proliferation induced by GLP1R agonists. In contrast, the loss of ACO2 activity and the resulting depletion of D-2HG were parallel metabolic consequences that were dispensable for the anti-proliferative response.

Collectively, our studies indicated that GLP1R agonism inhibited ACO2 and RNR at the functional rather than expression level, thereby disrupting the TCA cycle and nucleotide biosynthesis in mIDH glioma cells. The resulting dNTP depletion and block in DNA synthesis provide a metabolic basis for the anti-proliferative effect of GLP1R agonists in mIDH gliomas.

### GLP1R agonism induces NOS1-dependent NO production and S-nitrosylation of ACO2 and RNR

The coordinated loss of ACO2 and RNR activity in the absence of any change in their abundance pointed to a common post-translational mechanism. Although ACO2 and RNR function in distinct metabolic pathways and subcellular compartments, both are regulated by NO through S-nitrosylation of cysteine residues, and since NO diffuses freely across cellular membranes, a single NO source can modify targets in both the mitochondrial matrix and the cytosol^38,39^.

Treatment with LIR or NLY01 induced NO production in mIDH glioma cells in a GLP1R-dependent manner (Fig. 3A-3B). Immunoprecipitation of ACO2 or RRM1 followed by immunoblotting for S-nitrosocysteine revealed increased S-nitrosylation of both proteins after treatment with LIR or NLY01 in TS603 cells (Fig. 3C-3D). To further confirm these results, we coupled the biotin switch method to a proximity ligation assay (PLA), hereafter referred to as biotin-switch PLA, to probe for S-nitrosylation of ACO2 and RNR across all our mIDH glioma models^40,41^. Briefly, free thiols were blocked, S-nitrosothiols were then selectively reduced to free thiols with ascorbate, and the newly exposed thiols were labeled with the thiol-reactive biotinylating reagent HPDP-biotin^40^. S-nitrosylation of each target was then detected using an antibody against biotin together with an antibody against the target, followed by conjugation of species-specific PLA probes to complementary oligonucleotides^41^. The close proximity of the biotinylated cysteine residue and the target protein enabled oligonucleotide hybridization and ligation to generate a circular DNA template, which was then amplified to produce a fluorescence signal quantifiable by flow cytometry^41^. Compared with conventional biotin-switch Western blotting, the biotin-switch PLA approach allows sensitive, target-specific detection and quantification of protein S-nitrosylation at the single-cell level by flow cytometry^40,41^. First, we established the specificity of the biotin-switch PLA in LIR-treated cells, wherein the fluorescent signal is expected to be detectable. Omitting either primary antibody or replacing both with isotype-matched IgG reduced the fluorescent signal to the level seen in untreated cells (Supplementary Fig. S3A-S3D. The fluorescent signal was similarly lost when biotinylation or ascorbate was omitted (Supplementary Fig. S3A-S3D), confirming that the readout requires both probes and the complete switch chemistry^40,41^. Using the biotin-switch PLA method, we confirmed that both ACO2 and RRM1 were S-nitrosylated in LIR- or NLY01-treated cells across six patient-derived and murine mIDH glioma models (Fig. 3E-3F and Supplementary Fig. S3E-S3F). These results suggested that GLP1R agonists induced NO production and S-nitrosylation of ACO2 and RNR in mIDH glioma models.

**Figure 3.**
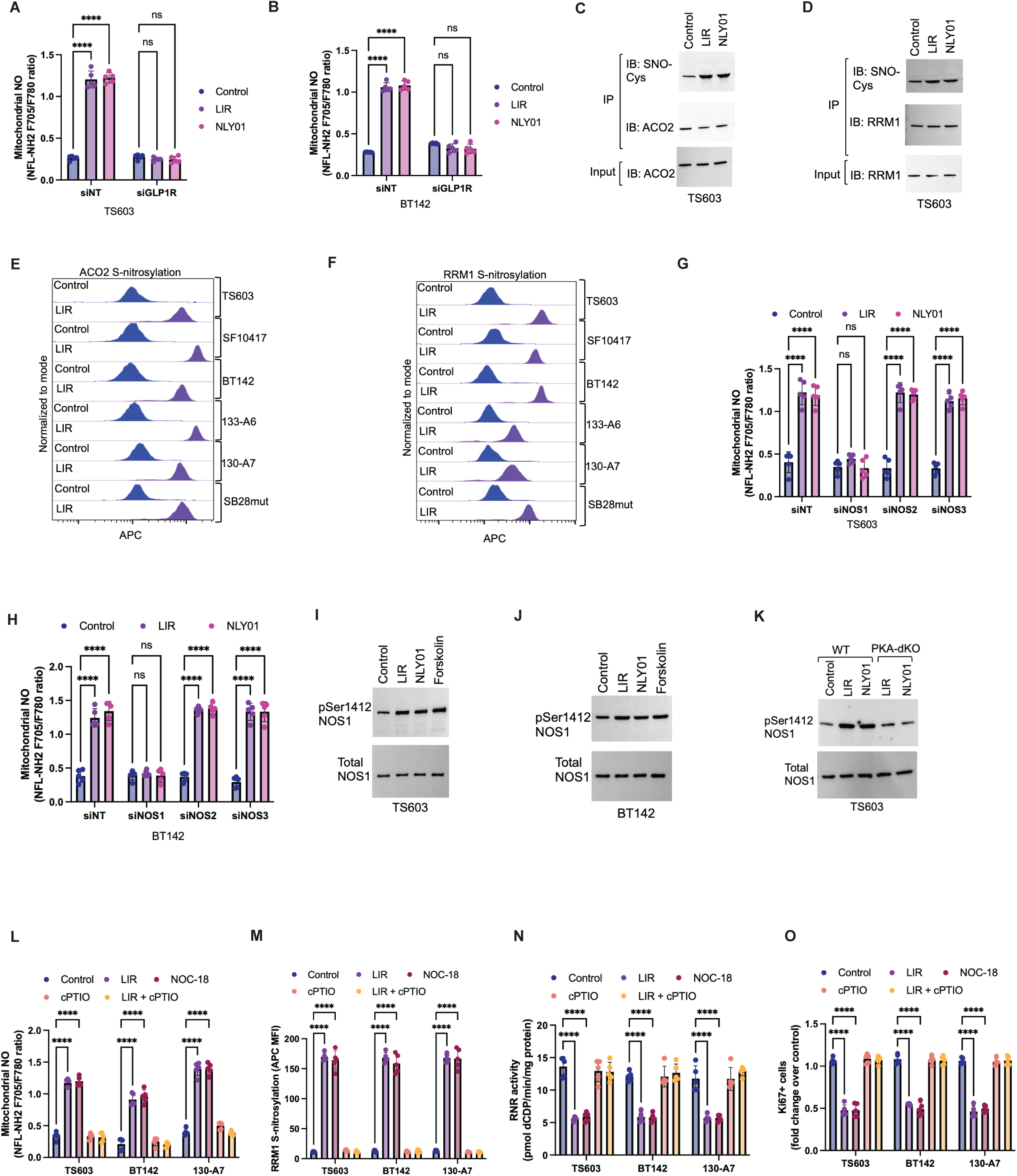
GLP1R agonism drives NOS1-dependent mitochondrial NO production and S-nitrosylation of ACO2 and RRM1. **(A, B)** Mitochondrial NO (NFL-NH2 F705/F780 ratio) in TS603 **(A)** and BT142 **(B)** transfected with siNT or siGLP1R and treated with vehicle, LIR or NLY01. **(C, D)** Immunoprecipitation of ACO2 **(C)** or RRM1 **(D)** from TS603 treated with vehicle, LIR or NLY01, followed by immunoblotting for S-nitrosocysteine (SNO-Cys). IP refers to the immunoprecipitate and IB to the immunoblot. **(E, F)** Representative biotin-switch proximity ligation assay traces showing S-nitrosylation of ACO2 **(E)** and RRM1 **(F)** in six mIDH glioma models. The traces show the LIR arm only; the NLY01 arm is quantified in Supplementary Fig. S3E and S3F. **(G, H)** Mitochondrial NO in TS603 **(G)** and BT142 **(H)** transfected with siNT, siNOS1, siNOS2 or siNOS3 and treated with vehicle, LIR or NLY01. **(I, J)** Immunoblot for NOS1 phosphorylated at Ser1412 and total NOS1 in TS603 **(I)** and BT142 **(J)** treated with vehicle, LIR, NLY01 or forskolin. **(K)** The same immunoblot in wild-type and PKA dKO TS603 cells. Wild-type cells were treated with vehicle, LIR or NLY01; PKA dKO cells were treated with LIR or NLY01. **(L-O)** TS603, BT142 and 130-A7 treated with vehicle, LIR, the NO donor NOC-18, cPTIO, or LIR together with cPTIO: mitochondrial NO **(L)**, RRM1 S-nitrosylation by biotin-switch PLA (APC MFI, **M**), RNR activity **(N)** and Ki67+ cells **(O)**. Flow cytometry histograms are normalized to mode and plotted on an identical axis range. Data are mean ± SD with individual biological replicates shown (n ≥ 3 per group). ****p<0.0001, ***p<0.001, **p<0.01, *p<0.05; ns, not significant.

To identify the enzyme driving NO production, we examined the effect of silencing NOS1, NOS2, or NOS3 in mIDH glioma cells (Supplementary Fig. S3G). Silencing NOS1 but not NOS2 or NOS3 abrogated NO production in TS603 and BT142 cells (Fig. 3G-3H). We then examined how GLP1R agonism activated NOS1. NOS1 can be phosphorylated by PKA on an activating serine residue (Ser1412). Treatment with a GLP1R agonist or the adenylyl cyclase activator forskolin induced NOS1 phosphorylation at Ser1412 in both TS603 and BT142 cells (Fig. 3I-3J). Importantly, the increase in NOS1 Ser1412 phosphorylation produced by LIR or NLY01 in wild-type cells was not seen in PKA double-knockout cells, while total NOS1 was unchanged (Fig. 3K). These results indicated that GLP1R agonism activated NOS1 through the cAMP-PKA pathway, placing NO production downstream of the cAMP-PKA step and completing the mechanistic chain from the GLP1R to NO production.

We next tested whether NO was sufficient to reproduce these modifications and their associated metabolic effects. Treatment with the NO donor diethylenetriamine NONOate^42^ (DETA NONOate; NOC-18) at a concentration producing intracellular NO comparable to that induced by LIR (Fig. 3L) increased S-nitrosylation of both ACO2 and RRM1 (Fig. 3M and Supplementary Fig. S3H) and reduced their enzymatic activities (Fig. 3N and Supplementary Fig. S3I), phenocopying the effects of GLP1R agonism. Conversely, scavenging NO with 2-(4-carboxyphenyl)-4,4,5,5-tetramethylimidazoline-1-oxyl-3-oxide (cPTIO)^43^ prevented LIR-induced S-nitrosylation of ACO2 and RNR and restored their enzymatic activities (Fig. 3L-3N and Supplementary Fig. S3H-S3I). Of note, NOC-18 phenocopied the anti-proliferative effect of LIR, whereas cPTIO rescued proliferation in LIR-treated cells (Fig. 3O). Collectively, these pharmacological and genetic experiments establish that GLP1R agonism activates cAMP-PKA signaling and induces NOS1-dependent NO production, leading to S-nitrosylation and inhibition of ACO2 and RNR, suppression of DNA synthesis, and reduced proliferation in mIDH glioma cells.

### VOR relieves D-2HG-mediated inhibition of PDH and generates mitochondrial superoxide

We recently demonstrated that mIDH glioma models differ in their intrinsic sensitivity to VOR^26^. Using a panel of patient-derived models, we showed that VOR inhibits clonogenic growth, with IC50 values of <0.3 µM in one group and >5 µM in another group^26^. Resistance in the latter group is intrinsic rather than acquired and reflects continued D-2HG synthesis by PHGDH despite mIDH inhibition, resulting in persistently elevated intracellular D-2HG during VOR treatment^26^. We therefore refer to these groups as VOR-sensitive and VOR-resistant^26^. Consistent with the delayed tumor growth observed in the clinical setting, VOR exerts a predominantly cytostatic effect in VOR-sensitive mIDH glioma models^26^. We confirmed that VOR inhibits the proliferation of the mIDH glioma models used in this study without inducing overt cell death, indicating that all models are VOR-sensitive (Fig. 4A and Supplementary Fig. S4A).

**Figure 4.**
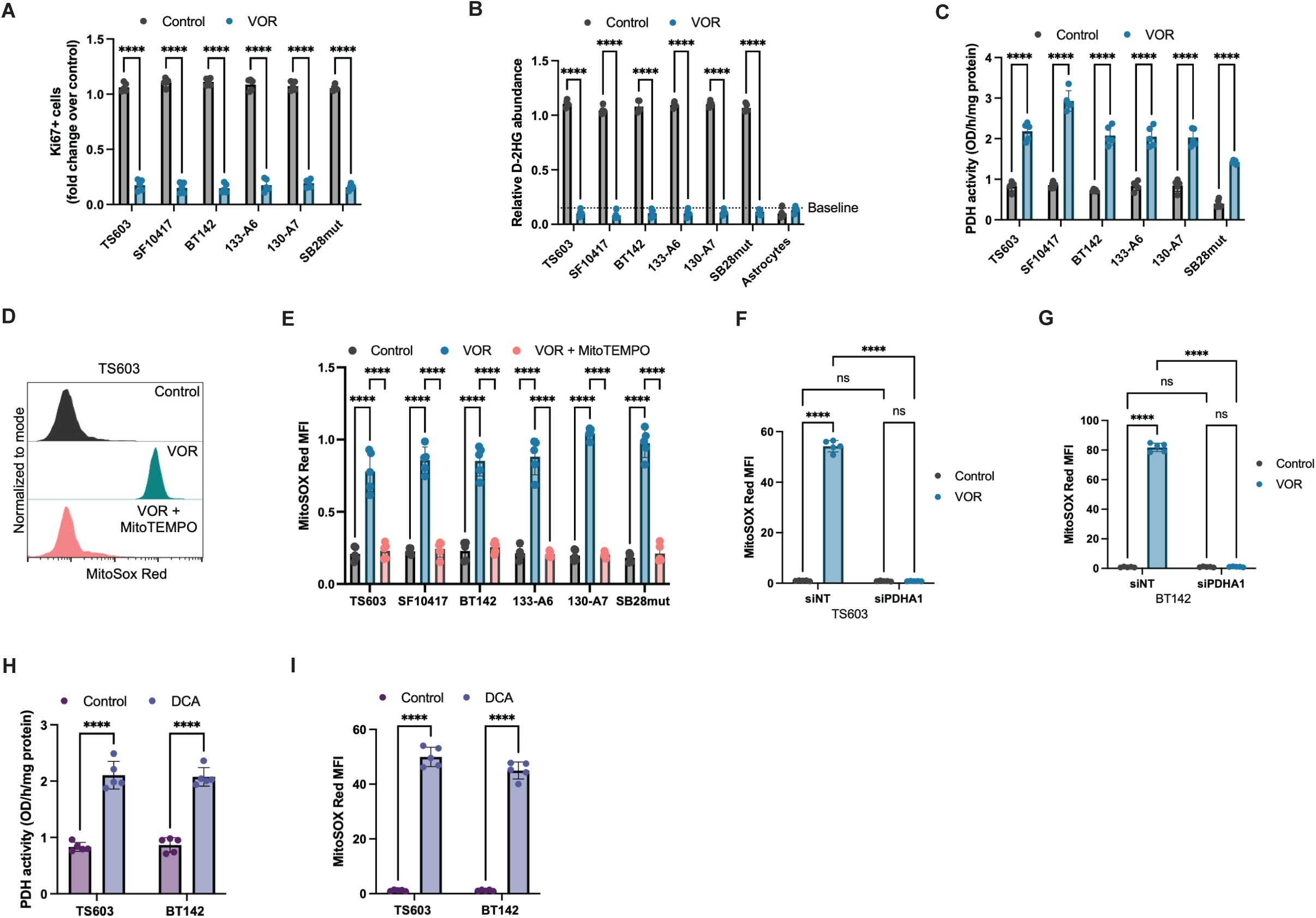
VOR relieves D-2HG-mediated suppression of PDH and generates PDH-dependent mitochondrial superoxide. **(A)** Ki67+ cells (fold change over control) in six mIDH glioma models treated with vehicle or VOR. **(B)** Relative intracellular D-2HG abundance in the same models, with normal human astrocytes shown for comparison. The dotted line marks the baseline defined as the D-2HG concentration in NHAs. **(C)** PDH activity (OD/h/mg protein) in the same models, vehicle versus VOR. **(D)** Representative MitoSOX Red labeling in TS603 treated with vehicle, VOR, or VOR with MitoTEMPO. **(E)** MitoSOX Red MFI across six models for the same three arms. **(F, G)** MitoSOX Red MFI in TS603 **(F)** and BT142 **(G)** transfected with siNT or PDHA1-directed siRNA and treated with vehicle or VOR. **(H)** PDH activity in TS603 and BT142 treated with vehicle or DCA. **(I)** MitoSOX Red MFI in the same cells and treatments. Flow cytometry histograms are normalized to mode and plotted on an identical axis range. Data are mean ± SD with individual biological replicates shown (n ≥ 3 per group). ****p<0.0001, ***p<0.001, **p<0.01, *p<0.05; ns, not significant.

In another study, we demonstrated that D-2HG suppressed pyruvate dehydrogenase (PDH) activity and reduced the oxidation of glucose-derived pyruvate via the TCA cycle in mIDH glioma cells^16^. However, it was unclear whether suppressing D-2HG production using VOR reversed this phenotype. VOR markedly reduced intracellular D-2HG in these models to levels comparable with those in normal human astrocytes (Fig. 4B), an effect that was accompanied by a significant increase in PDH activity (Fig. 4C).

We next asked whether restoration of PDH activity altered mitochondrial redox homeostasis. VOR increased MitoSOX Red labeling, indicating elevated mitochondrial superoxide levels, an effect that was reversed by MitoTEMPO, a mitochondria-targeted superoxide scavenger (Fig. 4D-4E). Silencing PDHA1 (Supplementary Fig. S4B) abolished VOR-induced superoxide production in TS603 and BT142 cells (Fig. 4F-4G), establishing a requirement for PDH in superoxide production. Conversely, pharmacological activation of PDH with dichloroacetate (DCA) phenocopied the effect of VOR and increased mitochondrial superoxide (Fig. 4H-4I). Collectively, these results indicate that VOR-mediated suppression of D-2HG restores PDH activity and drives PDH-dependent mitochondrial superoxide production in mIDH glioma cells.

### GLP1R agonist-induced NO and VOR-driven superoxide converge to generate peroxynitrite that is synergistically cytotoxic to mIDH glioma cells

Our findings above indicated that GLP1R agonists and VOR induce distinct alterations in mitochondrial redox by generating NO and superoxide, respectively. NO reacts rapidly with superoxide to form peroxynitrite (ONOO⁻), which can damage proteins, lipids, DNA, and mitochondria, leading to cell death^44–46^. We examined whether combining VOR with a GLP1R agonist triggered peroxynitrite production in mIDH glioma cells using a novel fluorescent probe that detects mitochondrial peroxynitrite^47^. While we did not observe peroxynitrite production in cells treated with LIR or NLY01 alone or VOR alone, combined treatment with VOR and either LIR or NLY01 markedly increased mitochondrial peroxynitrite across patient-derived and murine mIDH glioma models (Fig. 5A and Supplementary Fig. S5A). To determine whether both treatment-induced reactive species were required for peroxynitrite formation, we independently disrupted each pathway. Scavenging NO with cPTIO or mitochondrial superoxide with MitoTEMPO prevented peroxynitrite production in mIDH glioma cells treated with a combination of VOR and LIR (Fig. 5B). These findings demonstrated that GLP1R agonist-induced NO and VOR-induced superoxide were both required for peroxynitrite generation during combination therapy.

**Figure 5.**
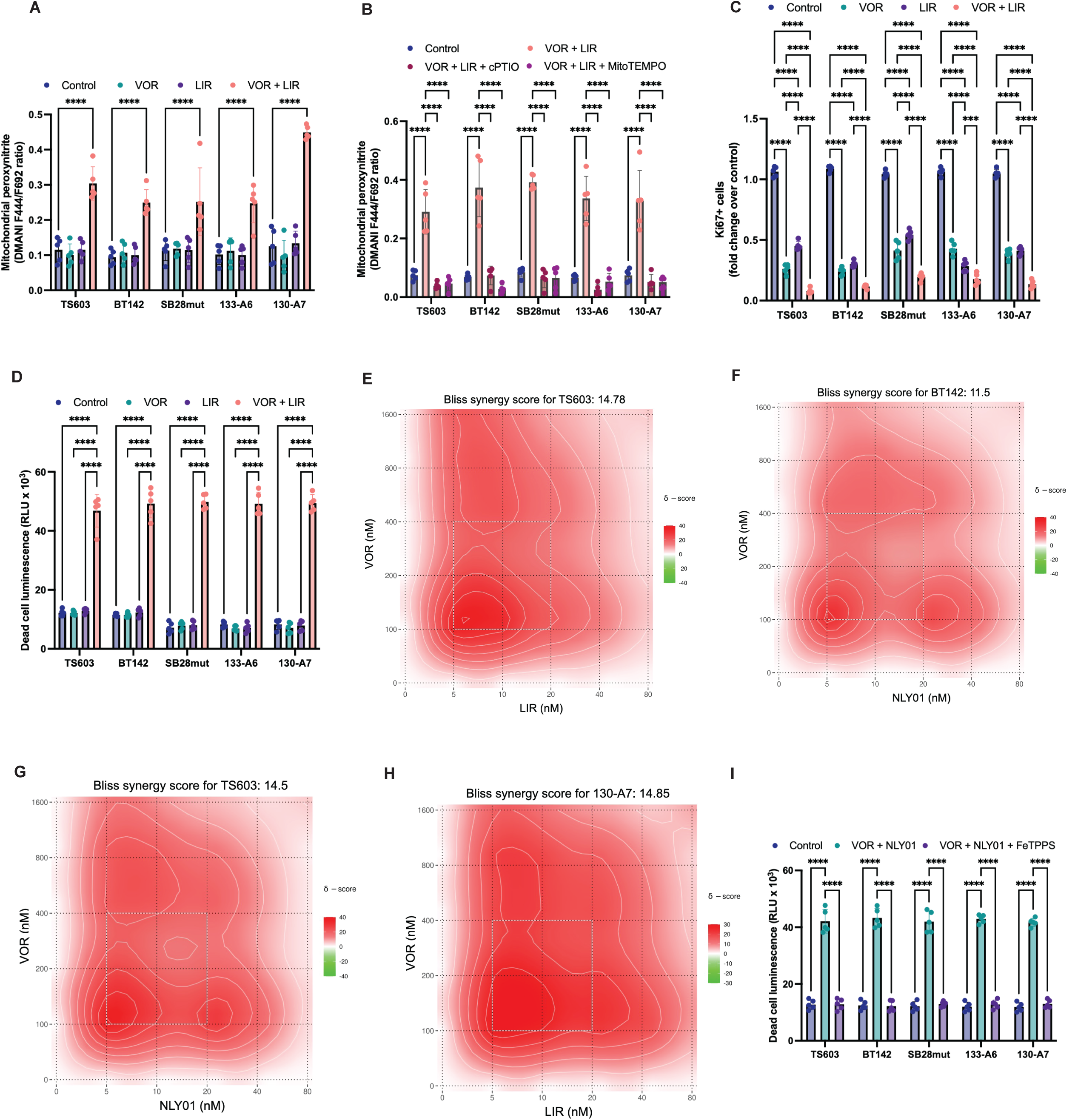
GLP1R agonist-derived NO and VOR-derived superoxide converge on peroxynitrite and produce synergistic cytotoxicity. **(A)** Mitochondrial peroxynitrite (DMANI F444/F692 ratio) in TS603, BT142, SB28mut, 133-A6 and 130-A7 treated with vehicle, VOR, LIR, or VOR with LIR. **(B)** Mitochondrial peroxynitrite in the same models treated with vehicle, VOR with LIR, or VOR with LIR plus either cPTIO or MitoTEMPO. **(C)** Ki67+ cells (fold change over control) in the same models and the arms shown in A. **(D)** Dead-cell luminescence (RLU x 10^3^) in the same models and arms. **(E-H)** Bliss synergy surfaces for the indicated agonist-VOR pairs, with the summary Bliss score above each panel: TS603 with LIR (E, 14.78), BT142 with NLY01 (F, 11.5), TS603 with NLY01 (G, 14.5) and 130-A7 with LIR (H, 14.85). Color encodes the delta score. **(I)** Dead-cell luminescence (RLU x 10^3^) in five mIDH glioma models treated with vehicle, VOR with NLY01, or VOR with NLY01 plus FeTPPS. Data are mean ± SD with individual biological replicates shown (n ≥ 3 per group). ****p<0.0001, ***p<0.001, **p<0.01, *p<0.05; ns, not significant.

We then examined the effect of combining VOR with a GLP1R agonist on tumor proliferation and cell death. Unlike monotherapy with a GLP1R agonist or VOR, which suppressed proliferation but did not induce cell death, combining LIR or NLY01 with VOR markedly increased cell death across mIDH glioma models (Fig. 5C-5D and Supplementary Fig. S5B). Dose-matrix analysis confirmed pharmacological synergy between VOR and LIR or NLY01 with Bliss synergy scores >10 in mIDH glioma models (Fig. 5E-5H).

We next determined whether peroxynitrite was necessary for this synergistic cytotoxicity. Treatment with the peroxynitrite decomposition catalyst FeTPPS (5,10,15,20-tetrakis(4-sulfonatophenyl) porphyrinatoiron(III) chloride) abrogated cell death induced by combined VOR and NLY01 treatment, implicating peroxynitrite as a mediator of the cytotoxic response^48,49^ (Fig. 5I). Collectively, these results indicate that GLP1R agonist-induced NO and VOR-induced superoxide converge to generate peroxynitrite, providing a mechanistic basis for the synergistic cytotoxicity of the combination.

### Peroxynitrite generated by the combination of VOR and a GLP1R agonist damages DNA and lipids, leading to apoptosis in mIDH glioma cells and intracranial tumors

Having established that peroxynitrite mediates the cytotoxicity induced by combining VOR with a GLP1R agonist, we sought to define the molecular mechanism underlying cell death. Peroxynitrite can induce irreversible or poorly reversible oxidative and nitrative damage to proteins, lipids, DNA, and mitochondrial components^44,50^. We confirmed that, unlike monotherapy, the combination of VOR and LIR uniquely induced lipid peroxidation in TS603 and BT142 cells (Fig. 6A-6B). This effect was rescued by the peroxynitrite decomposition catalyst FeTPPS, confirming that lipid peroxidation was dependent on peroxynitrite production.

**Figure 6.**
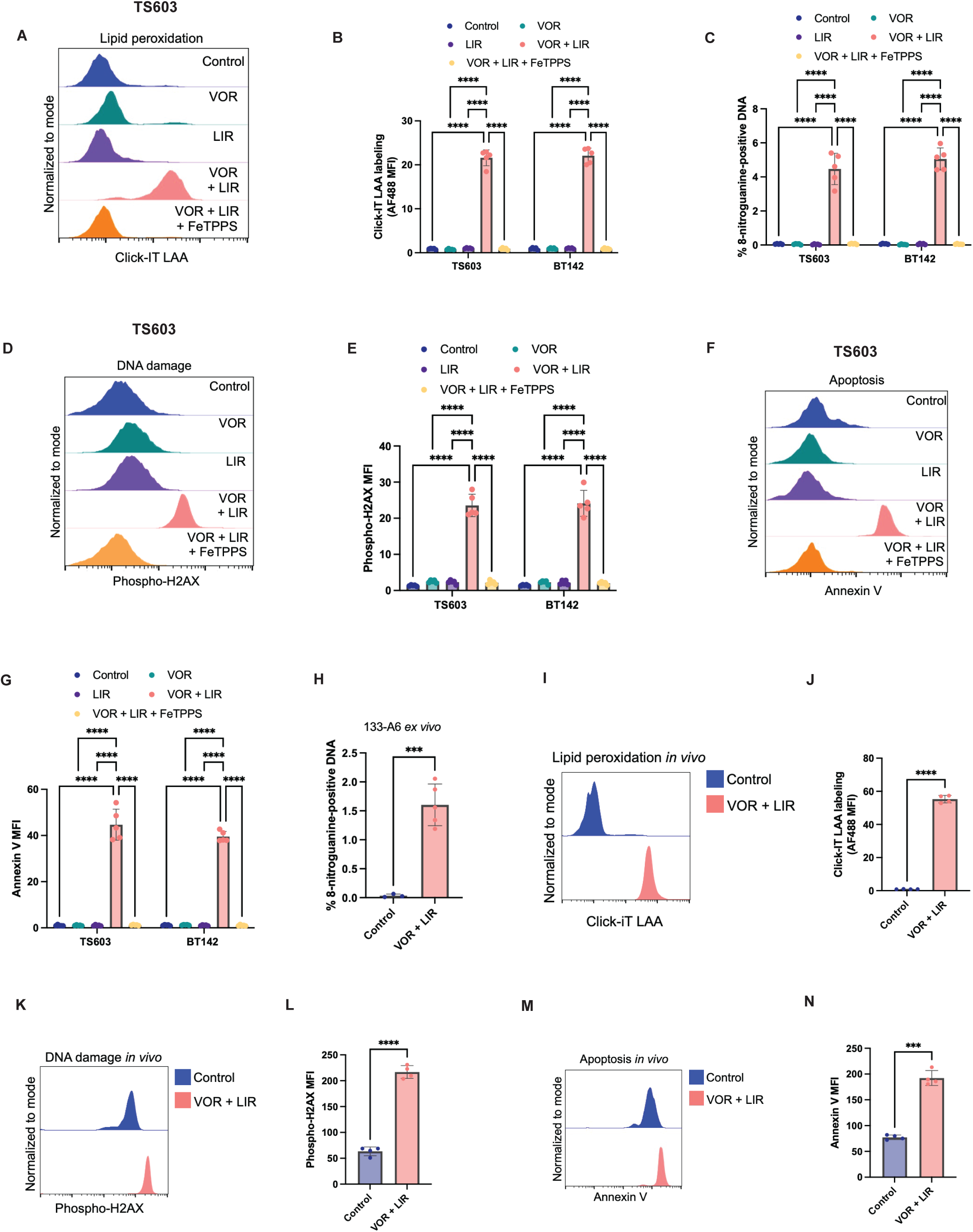
Peroxynitrite produced by the combination damages lipids and DNA and triggers apoptosis in mIDH glioma cells and intracranial tumors. **(A)** Representative Click-iT lipid peroxidation (linoleamide alkyne, LAA) labeling in TS603 treated with vehicle, VOR, LIR, VOR with LIR, or VOR with LIR plus FeTPPS. **(B)** Click-iT LAA labeling (AF488 MFI) in TS603 and BT142 across the same five arms. **(C)** 8-nitroguanine-positive DNA as a percentage of total, same cells and arms. **(D)** Representative phospho-H2AX labeling in TS603, same five arms. **(E)** Phospho-H2AX MFI in TS603 and BT142, same arms. **(F)** Representative annexin V labeling in TS603, same five arms. **(G)** Annexin V MFI in TS603 and BT142, same arms. **(H-N)** Mice bearing intracranial 133-A6 tumors treated with vehicle or VOR with LIR for approximately 7 days, then euthanized with tumor tissue resected and analyzed by flow cytometry. **(H)** 8-nitroguanine-positive DNA. **(I, J)** Representative Click-iT LAA labeling **(I)** and Click-iT LAA MFI **(J)**. **(K, L)** Representative phospho-H2AX labeling **(K)** and phospho-H2AX MFI **(L)**. **(M, N)** Representative annexin V labeling **(M)** and annexin V MFI **(N)**. Flow cytometry histograms are normalized to mode and plotted on an identical axis range. Data are mean ± SD with individual biological replicates shown (n ≥ 3 per group). ****p<0.0001, ***p<0.001, **p<0.01, *p<0.05; ns, not significant.

Because peroxynitrite can also nitrate nucleic acids, we next examined whether the combination induced nitrative DNA damage. Combined GLP1R agonism and VOR markedly increased the 8-nitroguanine content in DNA, whereas either agent alone had little effect (Fig. 6C), indicating that nitrative DNA damage is a unique readout of combination therapy. DNA damage as measured by phospho-H2AX abundance was markedly higher in cells treated with VOR and LIR relative to either agent alone or to untreated cells (Fig. 6D-6E). Concomitantly, apoptosis as measured by annexin V staining was higher in mIDH glioma cells treated with the combination of LIR and VOR (Fig. 6F-6G). FeTPPS reduced 8-nitroguanine accumulation and DNA damage and rescued apoptosis (Fig. 6C-6G).

To assess whether the combination of VOR and a GLP1R agonist recapitulated these effects in vivo, we examined mice bearing intracranial 133-A6 tumors treated with vehicle or the combination of VOR and LIR for approximately 7 days. Mice were then euthanized, and tumor tissue was resected for analysis by flow cytometry. As shown in Fig. 6H, nitrative DNA damage as measured by the 8-nitroguanine content in DNA was markedly higher in tumor tissue from mice treated with the combination of VOR and LIR relative to controls. Concomitantly, combination-treated tumors exhibited higher lipid peroxidation (Fig. 6I-6J), phospho-H2AX abundance (Fig. 6K-6L), and annexin V staining (Fig. 6M-6N) than controls. These results were consistent with our in vitro studies and indicated that combining a GLP1R agonist with VOR induced peroxynitrite production that caused oxidative damage to lipids and nitrative damage to DNA, leading to apoptotic cell death in mIDH glioma cells and intracranial tumors in vivo.

### The combination of a GLP1R agonist with VOR induces tumor regression and prolongs survival in intracranial mIDH glioma models

Next, we examined the therapeutic potential of combining a GLP1R agonist with VOR in mice bearing intracranial tumors in vivo. Mice were intracranially implanted with SB28mut cells and, once tumors were well established as measured by optical imaging (bioluminescence; BLI), mice were treated with vehicle, VOR, LIR, or the combination of VOR and LIR. Animals were monitored until they required euthanasia, in accordance with animal welfare guidelines. As shown in Fig. 7A, the combination of VOR and LIR significantly increased median overall survival relative to each monotherapy and to vehicle (5 days for vehicle-treated mice, 8 days for LIR-treated, 6 days for VOR-treated, and 13 days for the combination of VOR and LIR; p<0.0001). To quantify the anti-tumor effect in individual animals, we defined the best response for each mouse as the lowest post-treatment BLI signal relative to that animal’s pre-treatment signal. The BLI signal fell below the pre-treatment value in combination-treated mice, whereas tumors grew in the vehicle-, VOR-, and LIR-treated mice (Fig. 7B).

**Figure 7.**
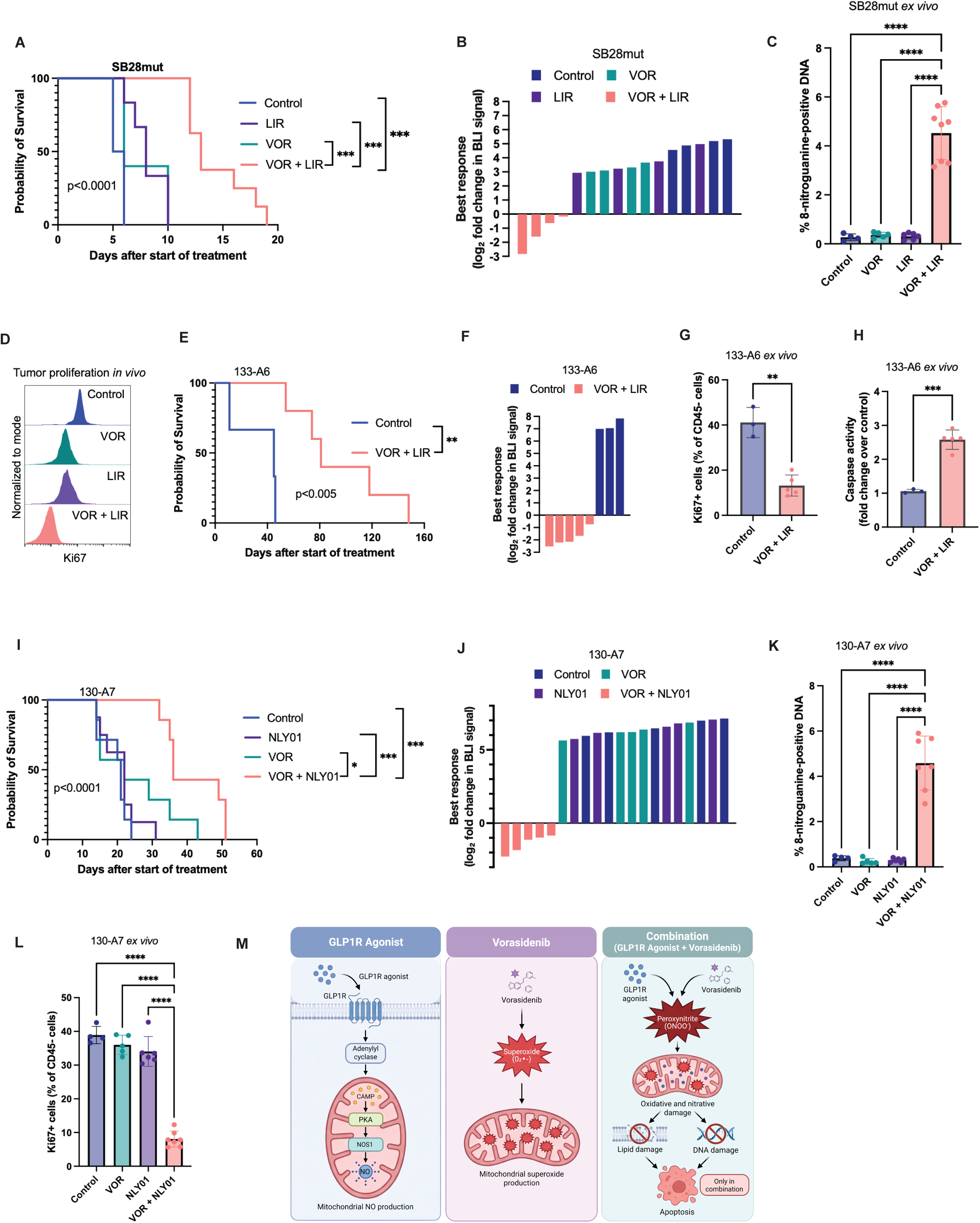
Combining a GLP1R agonist with VOR drives tumor regression and extends survival in three intracranial mIDH glioma models. **(A)** Kaplan-Meier survival of mice bearing intracranial SB28mut tumors treated with vehicle (Control), VOR, LIR, or the combination of VOR and LIR. Median overall survival was 5 days for vehicle, 8 days for LIR, 6 days for VOR and 13 days for the combination; p < 0.0001 by log-rank test (n=6 each for vehicle and LIR; n=5 for VOR; n=8 for the combination of VOR and LIR). **(B)** Best response of each SB28mut tumor, defined as its lowest post-treatment BLI signal relative to that animal’s own pre-treatment signal, expressed as a log2 fold change, ordered from lowest to highest and colored by treatment arm. The reference line at zero marks the pre-treatment value. n = 4 per arm. BLI fell below baseline in 4 of 4 combination-treated animals and in 0 of 12 vehicle-, VOR- and LIR-treated animals; p = 0.0005, Fisher’s exact test. **(C)** 8-Nitroguanine-positive DNA (%) in SB28mut tumor tissue resected at endpoint (n=4 for vehicle, n = 5 for VOR, n = 6 for LIR, n = 8 for the combination of VOR and LIR). **(D)** Representative flow cytometry histograms of Ki67 in CD45-negative cells from SB28mut tumor tissue resected at endpoint, for vehicle (Control), VOR, LIR and the combination. Quantification is shown in Supplementary Fig. S6. **(E)** Kaplan-Meier survival of mice bearing intracranial 133-A6 tumors treated with vehicle (Control) or the combination of VOR and LIR. Median overall survival was 45 days for vehicle and 81 days for the combination; p < 0.005 by log-rank test. **(F)** As in **(B)**, for 133-A6 tumors. n = 3 vehicle and n = 5 combination. BLI fell below baseline in 5 of 5 combination-treated animals and in 0 of 3 vehicle-treated animals; p = 0.018, Fisher’s exact test. **(G)** Ki67-positive cells (% of CD45-cells) and **(H)** caspase activity (fold change over control) in 133-A6 tumor tissue resected at endpoint. n = 3 vehicle and n = 5 combination. **(I)** Kaplan-Meier survival of mice bearing intracranial 130-A7 tumors treated with vehicle (Control), VOR, NLY01, or the combination of VOR and NLY01. Median overall survival was 21 days for vehicle, 22 days each for VOR and NLY01 and 36 days for the combination; p < 0.0001 by log-rank test. (n=7 each for vehicle, VOR and the combination; n=8 for NLY01). **(J)** As in **(B)**, for 130-A7 tumors. n = 5 per arm. BLI fell below baseline in 5 of 5 combination-treated animals and in 0 of 15 vehicle-, VOR- and NLY01-treated animals; p < 0.0001, Fisher’s exact test. **(K)** 8-Nitroguanine-positive DNA (%) and **(L)** Ki67-positive cells (% of CD45-cells) in 130-A7 tumor tissue resected at endpoint (n=4 for vehicle, n = 5 for VOR, n = 6 for NLY01, n = 7 for the combination of VOR and NLY01). **(M)** Schematic illustration of the mechanism of action of the combination. A GLP1R agonist drives cAMP-PKA signaling and NOS1-dependent mitochondrial NO production; VOR suppresses D-2HG, restores PDH activity and generates mitochondrial superoxide; in combination the two species form peroxynitrite, which damages lipids and DNA and triggers apoptosis. Created in BioRender. Panels B, F and J share a y-axis of -3 to +8 log2. Tumors were imaged before treatment and at scheduled intervals thereafter, with animals contributing one or two post-treatment measurements. Best response is the lowest available post-treatment measurement relative to that animal’s own baseline, rather than a nadir over a dense time course. Best response was determined in the subset of animals with a pre-treatment and at least one post-treatment BLI measurement. For SB28mut this comprised 4 of 6 vehicle, 4 of 6 LIR, 4 of 5 VOR and 4 of 8 combination-treated animals; for 130-A7 it comprised 5 of 7 vehicle, 5 of 7 VOR, 5 of 8 NLY01 and 5 of 7 combination-treated animals. All animals enrolled in the 133-A6 study contributed to panel F. Bars in C, G, H, K and L are mean +/- SD with individual animals shown as points. ****p<0.0001, ***p<0.001, **p<0.01, *p<0.05; ns, not significant.

Consistent with our in vitro studies, analysis of tumor tissue resected from mice at the endpoint showed a marked increase in the 8-nitroguanine content of DNA uniquely in mice treated with the combination of LIR and VOR (Fig. 7C). This molecular effect was accompanied by a significantly higher reduction in tumor cell proliferation as measured by the proportion of CD45-Ki67+ cells in combination-treated tumors relative to each monotherapy and to control (Fig. 7D and Supplementary Fig. S6).

To extend these results in a second model, we examined the effect of treatment with the combination of VOR-LIR on mice bearing intracranial 133-A6 tumors. Combination therapy significantly extended median overall survival (45 days for vehicle-treated mice vs. 81 days for mice treated with a combination of VOR and LIR; p<0.005; Fig. 7E). Similar to the SB28mut model, the BLI signal fell below the pre-treatment value in combination-treated mice, whereas vehicle-treated tumors showed growth (Fig. 7F). Tumor tissue resected from the combination-treated mice at the endpoint showed a marked reduction in Ki67+ cells and a concomitant increase in caspase activity relative to vehicle controls (Fig. 7G-7H).

Finally, we examined the effects of treatment with VOR, NLY01, or their combination on mice bearing intracranial 130-A7 tumors. In this model, median overall survival was unchanged by either monotherapy and was significantly extended only by the combination of VOR and NLY01 (21 days for vehicle, 22 days each for VOR and NLY01, and 36 days for the combination; p<0.0001; Fig. 7I). Like the SB28mut and 133-A6 models, BLI signal fell below the pre-treatment value in combination-treated mice, whereas vehicle-, VOR- and NLY01-treated mice showed tumor growth (Fig. 7J). Analysis of tumor tissue resected at endpoint showed that the combination of VOR and NLY01 uniquely increased 8-nitroguanine content in DNA (Fig. 7K) and reduced the fraction of CD45-Ki67+ cells (Fig. 7L) relative to the monotherapies and controls. Since suppression of growth cannot, by itself, reduce the BLI signal below its starting value, the responses in Fig. 7B, 7F, and 7J indicate that the combination induces a cytotoxic loss of viable tumor cells in all three models, although tumors ultimately progressed in every animal. Collectively, our studies indicate that the combination of a GLP1R agonist with VOR is synergistically cytotoxic to mIDH gliomas due to the production of lethal peroxynitrite (Fig. 7M) and provide proof of concept for the therapeutic potential of the combination in multiple intracranial mouse models of mIDH glioma.

## DISCUSSION

mIDH gliomas are primary brain tumors that predominantly affect younger adults, with a median age of approximately 40 years^1–3^. Although patients with mIDH gliomas survive longer than those with IDH wild-type glioblastoma, the disease remains incurable^1–3^. The approval of vorasidenib (VOR) as a first-line therapy for patients with grade 2 mIDH glioma represents a major advance^9–12^. However, while VOR increases progression-free survival and delays the time to the next intervention, patients eventually experience tumor progression^10–14^. There is an unmet need for combination strategies that convert growth suppression into tumor cell death. Here, using clinically relevant patient-derived and murine models, we identified and mechanistically validated GLP1R agonists as a clinically actionable drug class that synergistically induces lethality with VOR in mIDH gliomas. GLP1R agonism induced NOS1-dependent NO production, whereas VOR increased mitochondrial superoxide through restoration of PDH activity (Fig. 7M). When combined, these independently generated reactive species formed peroxynitrite, causing nitrative damage to DNA and lipids and triggering apoptotic cell death in vitro and in vivo Fig. 7M). Importantly, the combination induced tumor regression and prolonged survival relative to either monotherapy in mice bearing intracranial mIDH tumors. Together, these findings highlight the therapeutic potential of combining a GLP-1 receptor agonist with VOR in mIDH gliomas.

To the best of our knowledge, our study is the first to define a tumor-intrinsic role for GLP1R agonists in glioma. Both LIR and NLY01 activated cAMP-PKA signaling and induced NOS1-dependent NO production in mIDH glioma cells. NO suppressed proliferation by S-nitrosylation and inactivation of ACO2 and RNR, thereby disrupting TCA cycle activity and depleting the dNTP pools required for DNA synthesis, without inducing cell death. The ability of an NO donor to phenocopy these effects, and of an NO scavenger to reverse them, establishes NO as a causal mediator of the anti-proliferative response. These findings define a direct mechanistic link between GLP1R signaling and tumor cell metabolism that is independent of systemic, immune, or stromal effects.

The biological effects of NO in cancer are context-dependent^51,52^. Low, sustained NO can support proliferative and survival signaling, whereas higher concentrations induce cell-cycle arrest and nitrosative stress^51,52^. This framework may explain why chronic NOS2-derived NO from stromal or myeloid compartments has been associated with poor outcomes in several carcinomas^51,53^, while the acute, tumor-cell-intrinsic NO production induced here by GLP1R agonism is anti-proliferative. In mIDH glioma cells, GLP1R signals through PKA to NOS1, generating NO that inactivates ACO2 and RNR and suppresses proliferation. Although GLP1R-dependent activation of NOS has been described in vascular biology^54^, our findings extend this signaling axis to tumor cells and identify NOS1-derived NO as a direct effector of GLP1R agonism.

Our studies also identify a distinct role for VOR in mitochondrial redox regulation. We previously showed that D-2HG suppresses PDH activity and limits the entry of glucose-derived pyruvate into the TCA cycle^16^. We extended these results here to show that VOR restores PDH activity and increases mitochondrial superoxide production in mIDH gliomas. The requirement for PDHA1, together with the ability of pharmacological PDH activation to phenocopy this effect, links restoration of pyruvate oxidation to mitochondrial redox. These results indicate that metabolic normalization following mIDH inhibition is not just a pharmacodynamic consequence of drug-target engagement but simultaneously creates a targetable vulnerability.

Our work identifies a novel strategy to exploit therapeutically actionable redox vulnerabilities by pharmacologically generating complementary reactive species within tumor cells. Nitrosative chemistry has long been proposed as a strategy for inducing tumor cell death^55^, but its therapeutic application has been limited by the difficulty of generating sufficient NO within tumors without systemic toxicity. Our approach circumvents this limitation by pairing two agents that independently generate the precursors needed for peroxynitrite production. GLP1R agonism produces NO, whereas VOR-induced restoration of PDH activity generates mitochondrial superoxide. Their convergence therefore occurs selectively in cells in which both pharmacologic responses are engaged, namely GLP1R-expressing mIDH glioma cells undergoing effective mIDH inhibition. Consistent with this model, FeTPPS rescued cytotoxicity and attenuated the associated lipid peroxidation, 8-nitroguanine accumulation, and DNA damage response, placing peroxynitrite upstream of these lesions. These results are consistent with the established reactivity of peroxynitrite toward unsaturated lipids and guanine^44,46,50^. Although we cannot rule out the possibility that other reactive species contribute to 8-nitroguanine generation in our models, our studies indicate that combining a GLP1R agonist with VOR converts two independently generated reactive species into a third, more damaging species within the tumor cell, providing a mechanistic basis for the pharmacological synergy of the combination.

Our studies identify a broader strategy for designing drug combinations by pairing agents whose products react with one another, rather than by targeting the same or parallel signaling pathways. Under this framework, each drug generates a distinct reactive species whose interaction produces a cytotoxic species absent with either agent alone. This provides a potential means of overcoming a central limitation of mIDH inhibition, i.e., although VOR suppresses proliferation and slows tumor growth, its anti-tumor activity is predominantly cytostatic. Our findings show that this property can itself be exploited therapeutically. By restoring PDH activity, VOR generates mitochondrial superoxide, which serves as a substrate for peroxynitrite formation when GLP1R agonism supplies NO. The combination, therefore, does not simply intensify suppression of D-2HG signaling. Instead, it exploits a biochemical consequence of successful mIDH inhibition to create a cytotoxic mechanism that is absent with either treatment alone.

To the best of our knowledge, this study also provides the first evidence of the anti-tumor activity of a GLP1R agonist against an intracranial tumor and the first evaluation of a GLP1R agonist in combination with an mIDH inhibitor. Prior studies of GLP1R agonists in cancer have largely focused on obesity-associated tumors and have implicated growth-signaling pathways such as AMPK-mTOR or PI3K-AKT, as well as remodeling of the tumor microenvironment^56–62^. Our findings identify a distinct tumor-intrinsic mechanism in which GLP1R signaling drives NOS1-dependent NO production, thereby establishing nitrosative chemistry as a direct consequence of GLP1R activation. Importantly, NO becomes cytotoxic only in the presence of superoxide generated by mIDH inhibition. This tumor-intrinsic mechanism is supported by experiments using human patient-derived mIDH glioma cells in the absence of immune or stromal compartments, in which GLP1R silencing abolished the response.

Our studies provide a strong rationale for clinical evaluation of GLP1R agonists in combination with VOR in patients with mIDH glioma. An important translational feature of this combination strategy is that the mechanism of action defines a potential biomarker of sensitivity. The superoxide arm of the combination depends on restoration of PDH activity and, therefore, requires effective suppression of D-2HG by VOR. This distinction may be clinically important because our recent studies indicate that a subset of mIDH gliomas is intrinsically resistant to VOR, maintaining D-2HG production despite mIDH inhibition via a phosphoglycerate dehydrogenase (PHGDH)-driven metabolic bypass mechanism^26^. This distinction is important because tumors can exhibit a substantial reduction in D-2HG after VOR treatment yet remain biologically resistant if residual D-2HG synthesis exceeds the threshold required to maintain the mIDH state^26^. Thus, the relevant biomarker is not simply a reduction in D-2HG, but rather a sufficiently complete suppression of D-2HG production to permit restoration of PDH activity and the generation of the superoxide required for combination activity.

The mechanistic framework above highlights the opportunity for metabolic imaging to guide patient stratification. Conventional ^1^H-magnetic resonance spectroscopy (^1^H-MRS) measures steady-state D-2HG abundance, but its sensitivity is unlikely to distinguish a truly responsive tumor, in which D-2HG falls to near-baseline levels, from an intrinsically resistant tumor that retains a small but biologically meaningful residual D-2HG pool (less than 1 mM, and therefore likely below the detection threshold of ^1^H-MRS^63,64^). Both VOR-sensitive and resistant tumors may, therefore, show an apparent treatment-associated decrease in ^1^H-MRS-detectable D-2HG. In contrast, we recently demonstrated that deuterium metabolic imaging using deuterium-labeled α-ketoglutarate directly interrogates ongoing D-2HG synthesis and can distinguish tumors in which VOR has effectively extinguished D-2HG production from those in which residual synthesis persists despite treatment^26^. Together, these findings support biomarker-guided clinical development of the combination, with deuterium metabolic imaging used to identify effective D-2HG suppression and tumor pharmacodynamic markers used to confirm GLP1R engagement.

A key translational question is whether clinically achievable GLP1R agonist exposure will be sufficient to reproduce the cytotoxicity observed in our preclinical models. Our in vivo studies demonstrate therapeutic efficacy at low doses of GLP1R agonists. We administered both agents intranasally, a route that provides direct nose-to-brain access^65,66^, to avoid systemic body weight loss and increase brain exposure. Both LIR and NLY01 were administered at 0.4 mg/kg/day, which, by body-surface-area scaling, corresponds to approximately 2.3 mg per dose in a 70-kg adult. For LIR, which is also dosed daily in patients, this falls within the 1.8 to 3.0 mg daily range approved for chronic use in type 2 diabetes and obesity. Since LIR was administered intranasally, systemic exposure was likely a fraction of that achieved by subcutaneous dosing at the same dose, so the anti-tumor effect of LIR was observed at or below an exposure already achieved chronically in patients^67,68^. For NLY01, although the comparison is less direct because our mice were dosed daily while patients received weekly dosing, the per-dose equivalent of approximately 2.3 mg in our mice is below the lowest dose evaluated in the phase 2 Parkinson’s Disease trial, whereas the corresponding weekly cumulative exposure is approximately 2.7-fold above the highest dose evaluated in that trial^34^. Of note, these doses are equivalent by mass but not by molarity. NLY01 has a molecular weight of 54,290 Da, compared with 3,751 Da for LIR, so 0.4 mg delivers 7.37 nmol of NLY01, compared with 106.6 nmol of LIR, an approximately 14-fold lower molar dose. Given that the IC50 is comparable for LIR and NLY01 in vitro, the in vivo efficacy of NLY01 at a substantially lower molar dose most likely reflects its longer half-life rather than greater intrinsic potency. Of note, administering either GLP1R agonist in combination with VOR resulted in pharmacodynamic evidence of drug-target engagement, as shown by DNA nitration, lipid peroxidation, and activation of a DNA damage response and apoptosis in tumor tissue, effects that were accompanied by tumor regression and prolonged survival in mice bearing intracranial mIDH gliomas. The observation that two structurally distinct GLP1R agonists converge on the same mechanism and outcome supports GLP1R agonism as the relevant therapeutic principle rather than any single agent.

It should be noted that VOR was specifically developed as a brain-penetrant mIDH inhibitor, whereas brain penetrance varies substantially among GLP1R agonists. Although NLY01 was developed specifically for neurological applications^34^, direct measurements of drug concentration within human brain tissue are lacking. This issue may be particularly relevant in non-enhancing mIDH gliomas, where the blood-brain barrier is relatively intact. The remaining translational challenge is therefore not proof of intracranial activity but the selection of the optimal dose and route of administration that will provide clinically meaningful antitumor activity in combination with VOR. Perioperative window-of-opportunity studies will therefore be important for establishing whether systemic dosing achieves comparable intratumoral exposure in patients.

Our studies have some limitations that warrant consideration. We established GLP1R dependence in vitro but not in vivo. Confirming GLP1R dependence in vivo using mIDH glioma models with genetically silenced GLP1R will strengthen our mechanistic framework. Another limitation is that both agonists were delivered intranasally in our in vivo studies. Further studies comparing subcutaneous vs. intranasal administration in mice bearing intracranial mIDH gliomas are needed to assess whether systemic dosing is efficacious in vivo. Our efficacy studies included hosts with varying immune statuses, which makes it difficult to assess the contribution of the tumor immune microenvironment to the anti-tumor efficacy of the combination therapy. Further studies are needed to understand the effects of combining a GLP1R agonist with VOR on anti-tumor immunity and response to immunotherapy.

In summary, our studies identify a therapeutically actionable strategy for converting the cytostatic effects of mIDH inhibition and GLP1R agonism into synergistic tumor cell killing. Suppression of D-2HG by VOR restores PDH activity and increases mitochondrial superoxide, which reacts with GLP1R agonist-induced NO to generate cytotoxic peroxynitrite. Thus, the metabolic changes triggered by mIDH inhibition create a vulnerability that can be exploited by GLP1R agonism to drive tumor cell death. This mechanism is supported by pharmacological, genetic, biochemical, and in vivo evidence and translates into tumor regression and prolonged survival in intracranial mouse models of mIDH glioma. Taken together, our findings establish redox stress induced by targeted therapies as an exploitable vulnerability for converting cytostatic tumor control into tumor cell death.

## MATERIALS AND METHODS

### Study design

This study assessed the therapeutic potential of combining a GLP1R agonist with VOR in preclinical mIDH glioma models. Clinically relevant patient-derived and genetically engineered murine mIDH glioma models were used to mechanistically interrogate the effects of each therapy alone and in combination^26,27^. Sample sizes were chosen based on effect sizes from prior studies. The samples represent biological replicates, defined as independently cultured and/or treated populations for in vitro studies and independent animals for in vivo studies. No statistical method was used to predetermine sample size. No data points were excluded unless justified because of technical errors. Animals without a post-treatment BLI measurement were not included in the best-response analysis. All enrolled animals were included in the survival analyses. Studies were performed in accordance with the National Institutes of Health Guide for the Care and Use of Laboratory Animals, and study protocols were approved by the University of California, San Francisco (UCSF) Institutional Animal Care and Use Committee (IACUC; protocol # AN209338). Mice were randomly assigned to treatment groups, and investigators were not blinded during data collection and analysis. Study endpoints were in accordance with IACUC guidelines.

### Statistical analysis

All experiments were performed with at least three independent biological replicates (n≥3), defined as independently cultured and treated cell populations for in vitro studies and independent animals for in vivo studies. Results are expressed as mean ± SD as indicated in figure legends. Statistical analyses were performed in GraphPad Prism 10. Sample sizes were determined based on prior studies and feasibility considerations. No formal power calculation was performed. Two groups were compared using unpaired two-tailed Welch’s t tests; three or more groups were analyzed by one-way ANOVA; and multi-factorial comparisons were analyzed by two-way ANOVA. Multiple comparisons were corrected using Tukey’s, Šídák’s, or Dunnett’s tests as appropriate. The normality of data distribution was checked using the Shapiro-Wilk, D’Agostino-Pearson omnibus, Anderson-Darling, or the Kolmogorov-Smirnov (distance) method. Mouse survival curves were compared using log-rank Mantel-Cox tests. The significance threshold was α=0.05 (two-sided, normal-based 95% confidence interval).

### Cell culture

All mIDH glioma models used in this manuscript have previously been described^26,27,69–72^. TS603 (RRID: CVCL_A5HW) was obtained from Dr. Timothy Chan (Cleveland Clinic); SF10417 (no RRID) from Dr. Joseph Costello (University of California, San Francisco); BT142 (RRID: CVCL_D718) from ATCC; 133-A6 and 130-A7 (RRID unknown) cells were provided by Dr. Timothy Phoenix (University of Cincinnati), and SB28mut (RRID unknown) by Dr. Hideho Okada (University of California, San Francisco). These cell lines are not distributed by the authors and must be requested directly from these investigators. TS603 was derived from a patient with a grade 3 anaplastic oligodendroglioma and carries an endogenous heterozygous IDH1 R132H mutation together with 1p/19q codeletion^69^. SF10417 is a patient-derived line established from a recurrent, temozolomide-treated IDH1 R132H-mutant oligodendroglioma with a 1p/19q codeletion^70^. BT142 was established from a grade 3 IDH1 R132H-mutant astrocytoma^71^. The 133-A6 and 130-A7 models are female murine mIDH glioma lines derived from autochthonous tumors generated by in utero electroporation, in which IDH1 R132H and PIK3CA E545K were co-expressed, with loss of Cic and Fubp1^27^. 133-A6 carries a Cdkn2a deletion and gives rise to fully penetrant tumors that histologically resemble human grade 2-3 IDH-mutant, 1p/19q-codeleted oligodendroglioma^27^. In 130-A7, Trp53 deletion replaces Cdkn2a loss, thereby shortening latency and shifting histology toward high-grade astrocytoma^27^. The two lines, therefore, model the two histological subtypes of mIDH glioma. SB28mut is a syngeneic murine line derived from the SB28 GBM model and engineered to carry the IDH1 R132H mutation^72^.

mIDH glioma cells were maintained as neurospheres in serum-free Neurobasal-A medium supplemented with B27 and N2 supplements, 2 mM glutamine, 20 ng/mL each of EGF, FGF, and PDGF, and antibiotics. Normal human astrocytes (NHA) and normal human oligodendrocytes (NHO) were used as non-neoplastic controls. HEK293 cells stably expressing GLP1R and isogenic GLP1R-knockout HEK293 cells were maintained in Dulbecco’s modified Eagle’s medium (DMEM) with 2 mM glutamine and antibiotics and were used to establish the specificity of LUXendin645 labeling. All patient-derived glioma models were authenticated by short tandem repeat profiling (Cell Line Genetics) and tested for mycoplasma contamination upon receipt and at 6-month intervals thereafter. Experiments were performed using cultures maintained for no more than 30 passages following thawing.

### Drug treatments

All inhibitors were obtained from MedChemExpress or Sun-shine Chemicals unless otherwise specified. Unless otherwise indicated, cells were treated for 72 h prior to analysis. This time point was elected to permit downstream biological effects of D-2HG suppression to develop while minimizing secondary adaptations associated with prolonged culture. Cells were treated with vehicle (DMSO) or 1 uM VOR for 72 h. The 1 uM concentration was selected because it produced near-maximal suppression of D-2HG in VOR-sensitive models while remaining within the range of intratumoral VOR concentrations (∼1.5-7 uM) measured in tissue resected from patients with mIDH glioma in the INDIGO trial^9^. The GLP1R agonists LIR, LIX, and NLY01 were used at 20 nM for 72 h. Across the six mIDH glioma models, the IC50 was 7.4 to 19.2 nM for LIR, 10.1 to 18.2 nM for NLY01, and 8.9 to 15.0 nM for LIX (Fig. 1E-1G), placing 20 nM at 1.0 to 2.7 times the IC50 in every agonist and model combination tested. Because all treatments were performed in serum-free medium, the nominal concentration approximates the free concentration available to the cells. A single concentration was used across the three agonists for mechanistic comparability rather than to reproduce the plasma exposure of any individual agent. The GLP1R antagonist exendin-(9-39) (EXE) was used at 1 µM for 24 h before treatment with the GLP1R agonist. The adenylyl cyclase activator forskolin was used at 10 µM, and the adenylyl cyclase inhibitor SQ22536 at 100 µM. The nitric oxide donor DETA NONOate (NOC-18) was used at 100 µM, and the nitric oxide scavenger cPTIO at 10 µM. The peroxynitrite decomposition catalyst FeTPPS was used at 10 µM, with cells pretreated for 1 h before the experimental treatment and FeTPPS maintained throughout the subsequent treatment. The mitochondria-targeted superoxide scavenger MitoTEMPO was used at 10 µM, with a 1-hour pretreatment followed by continued exposure for 24 hours. The pyruvate dehydrogenase kinase inhibitor dichloroacetate (DCA) was used at 10 mM. For α-ketoglutarate rescue, cells were co-treated with cell-permeable octyl-α-ketoglutarate at 1 mM. For deoxyribonucleoside rescue, the medium was supplemented with 100 µM each of 2′-deoxyadenosine, 2′-deoxycytidine, 2′-deoxyguanosine, and thymidine for the duration of treatment.

### IC50 and synergy

Cells were treated with vehicle or a 10-point serial dilution of drug. The % of Ki67-positive cells were measured by flow cytometry after fixation and permeabilization or using the Lumit hKi67 assay (Promega) according to the manufacturer’s instructions. Cells were treated with a matrix of concentrations of VOR and either LIR or NLY01 for 72 h. The % of Ki67-psotive cells were measured as described above. Bliss synergy scores were calculated using SynergyFinder v3. Scores greater than 10 were considered to indicate synergy.

### GLP1R staining

Cells were dissociated with Accutase, washed in HBSS containing 1% BSA, and incubated with LUXendin645 at 250 nM for 60 minutes at 37°C in the dark. Cells were washed twice and analyzed by flow cytometry on a MACSQuant 10 analyzer (RRID: SCR_020268) in the R1/APC channel using 635-nm excitation and a 655-730-nm emission filter. Unstained cells processed in parallel defined the background, and labeling is reported as background-subtracted median fluorescence intensity. Specificity was established in HEK293 cells expressing GLP1R and in isogenic GLP1R-knockout HEK293 cells processed in the same experiment.

### DNA synthesis

DNA synthesis DNA synthesis was measured by EdU incorporation. Cells were pulsed with 10 µM EdU for 2 h at 37°C, harvested, fixed for 15 min at room temperature, and permeabilized in the saponin-based permeabilization buffer supplied with the kit for 15 min. Incorporated EdU was detected by copper-catalyzed click chemistry using the Click-iT Plus EdU Alexa Fluor 647 Flow Cytometry Assay Kit (Thermo Fisher Scientific, C10635) for 30 min at room temperature protected from light, according to the manufacturer’s instructions, followed by flow cytometry. Proliferation was additionally assessed by viable-cell counting using trypan-blue exclusion in parallel cultures at the indicated time points. EdU-positive cells were quantified among viable singlets. The same EdU pulse duration was maintained across conditions, including treatments that altered proliferation or S-phase duration.

### cAMP and PKA activity

Cells were equilibrated in assay medium containing 500 µM 3-isobutyl-1-methylxanthine (IBMX) for 10 min at 37°C and then stimulated with the indicated agonist for 15 min at 37°C. Reactions were terminated by aspiration of the medium and immediate addition of lysis buffer. Intracellular cAMP was quantified using the cAMP Complete ELISA Kit (Abcam, ab133051) according to the manufacturer’s instructions. Forskolin (10 µM for 15 min) was included as a positive control for adenylyl-cyclase activation.

PKA activity was measured in clarified cell lysates using the PKA Kinase Activity Assay Kit (Abcam, ab139435). PKA-dependent phosphorylation of a plate-bound peptide substrate was detected using a phospho-specific antibody, and absorbance was read at 450 nm. cAMP values were normalized to viable-cell number in replicate wells, whereas PKA activity was normalized to the protein mass loaded into each reaction.

### NO, superoxide, and peroxynitrite assays

Mitochondrial nitric oxide was measured using the mitochondria-targeted near-infrared ratiometric probe NFL-NH_2_. Cells were loaded with 10 µM NFL-NH₂ for 30 min at 37°C, washed twice with phenol-red-free HBSS, and incubated in fresh HBSS for 15 min before analysis on a spectral fluorescence plate reader. Fluorescence was recorded with approximately 650-nm excitation and emission collected at 705 and 780 nm; mitochondrial NO was reported as the F705/F780 ratio. DEA NONOate (100 µM for 30 min) was used as an NO-donor positive control, and parallel samples pretreated with 10 µM 7-nitroindazole for 30 min were used to test NOS1 dependence. Mitochondrial superoxide was measured using MitoSOX Red at 5 µM for 10 min at 37°C protected from light. Cells were washed twice with HBSS and analyzed by flow cytometry using 488- or 510-nm excitation and emission centered at 580 nm. Antimycin A (10 µM for 30 min) was included as a positive control. Mitochondrial peroxynitrite was measured using DMANI, a mitochondria-targeted ratiometric fluorescent probe for peroxynitrite^47^. Cells were loaded with DMANI at 37°C, washed with PBS, and analyzed on a spectrally compatible plate reader. Fluorescence was collected in the blue and red emission channels specified for DMANI, and mitochondrial peroxynitrite was quantified as the blue-to-red fluorescence ratio. Because the readout is ratiometric, untreated cells have a non-zero baseline ratio; values were expressed as fold change relative to untreated cells.

### S-nitrosylation assay

S-nitrosylated cysteines were labeled using the biotin-switch method. Cells were lysed in HEN buffer containing 250 mM HEPES, pH 7.7, 1 mM EDTA, and 0.1 mM neocuproine, supplemented with 2.5% SDS and 20 mM methyl methanethiosulfonate (MMTS). Free thiols were blocked for 30 min at 50°C with intermittent vortexing. Excess MMTS was removed by precipitation with four volumes of ice-cold acetone for 20 min at −20°C, and protein pellets were recovered by centrifugation and resuspended in HENS buffer containing 1% SDS. S-nitrosothiols were selectively reduced with 20 mM sodium ascorbate, and the nascent thiols were labeled with 1 mM HPDP-biotin for 1 h at room temperature with gentle mixing. Biotinylated proteins were captured on streptavidin-agarose resin for 1 h at room temperature or overnight at 4°C, washed extensively, eluted in reducing Laemmli buffer, and immunoblotted for ACO2 and RRM1. For the biotin-switch proximity-ligation assay, fixed and permeabilized cells were subjected to the corresponding in situ switch chemistry and incubated with a mouse monoclonal anti-biotin antibody together with rabbit antibodies against ACO2 or RRM1. Bound primary antibodies were detected using Duolink In Situ PLA Probe anti-Mouse PLUS and anti-Rabbit MINUS, followed by ligation and rolling-circle amplification using the Duolink In Situ Detection Reagents according to the manufacturer’s instructions. The PLA signal was quantified by flow cytometry on an MACSQuant 10 analyzer.

### ACO2 activity

ACO2 activity was measured using the Aconitase Assay Kit (Abcam, ab109712). Fresh cell pellets were extracted in the supplied aconitase-preservation solution and clarified by centrifugation. Aconitase-dependent formation of cis-aconitate from isocitrate was monitored kinetically at 240 nm every 20 s for 30 min at room temperature, and activity was calculated from the linear rate using the cis-aconitate extinction coefficient supplied with the kit. Samples were maintained on ice and assayed promptly because aconitase activity is labile.

### PDH activity

PDH activity was measured using the PDH Enzyme Activity Microplate Assay Kit (Abcam, ab109902). PDH was immunocaptured from detergent extracts for 3 h at room temperature, after which the NADH-coupled reaction was monitored kinetically at 450 nm for 30 min at room temperature. Activity was calculated from the linear increase in absorbance after subtraction of the blank.

### QPCR

Gene expression was measured by QPCR on a QuantStudio 5 system (ThermoFisher Scientific, RRID: SCR_018717), and data were normalized to β-actin. Total RNA was extracted using the TRIzol reagent and converted to cDNA using the High-Capacity cDNA Reverse Transcription Kit (Applied Biosystems/ThermoFisher Scientific, Cat# 4368814, RRID: AB_2801667). QPCR was performed using the TaqMan Fast Advanced Master Mix. Relative expression was calculated by the comparative Ct method, with ACTB as the reference transcript. (GLP1R: Hs01006332_m1; NOS1: Hs00167223_m1; NOS2: Hs01075529_m1; NOS3: Hs01574665_m1; PDHA1: Hs01049345_g1; ACTB: Hs01060665_g1; ACO2, Hs00426616_m1; RRM1: Hs01040698_m1).

### Gene silencing

GLP1R, NOS1, NOS2, NOS3, and PDHA1 were transiently silenced using ON-TARGETplus SMARTpool siRNA (Dharmacon/Horizon Discovery), with an ON-TARGETplus non-targeting pool used as the control. Cells were plated in antibiotic-free medium and transfected the following day at approximately 50-70% confluence with 25 nM total siRNA using DharmaFECT 1 according to the manufacturer’s instructions. Medium was replaced after 16–24 h. Knockdown was evaluated by QPCR at 48 h, and protein expression and functional endpoints were evaluated at 72 h unless otherwise indicated.

### Immunoblotting

Lysates or immunoprecipitated proteins were prepared as described above. Total cellular protein (∼20 µg) was separated on a 4-20% polyacrylamide gel (Bio-Rad) by sodium dodecyl sulfate-polyacrylamide gel electrophoresis and transferred onto Immobilon-FL PVDF membrane (Millipore). Membranes were blocked overnight in 5% milk (Santa Cruz Biotechnology) in TBST (20 mM Tris-HCl, pH 7.5, 500 mM NaCl, 0.1% Tween 20) at 4°C. Membranes were then washed 3 times for 5 min each in TBST and incubated with primary antibodies diluted in TBST for 1 h at room temperature. Following 3 washes of 10 min each with TBST, fluorescence-conjugated secondary antibodies were added for 1 h in TBST at room temperature. Membranes were washed thrice in TBST for 10 min each and developed using an iBright imager (ThermoFisher Scientific).

### Lipid peroxidation

Lipid peroxidation was measured using the Click-iT Lipid Peroxidation Imaging Kit with linoleamide alkyne (LAA; Thermo Fisher Scientific, C10445). Cells were incubated with 50 µM LAA during the final 2 h of treatment, washed with PBS, and fixed with 4% paraformaldehyde for 15 min at room temperature. Cells were permeabilized with 0.1% Triton X-100 for 10 min, and protein adducts generated by LAA-derived electrophilic aldehydes were detected by click reaction with the supplied Alexa Fluor 488 azide for 30 min at room temperature protected from light. After washing, fluorescence was measured by flow cytometry. Cells exposed to 100 µM cumene hydroperoxide for 2 h in the presence of LAA were included as a positive control. An LAA-omission control and a click-reagent-omission control were included to define nonspecific fluorescence. Results were reported as median Alexa Fluor 488 fluorescence among viable singlets and normalized to viable-cell number.

### Nitrative DNA damage

8-Nitroguanine was measured in fixed cells rather than purified genomic DNA because the modified base undergoes spontaneous depurination and can be lost during DNA isolation. Cells were harvested directly into ice-cold PBS, fixed within 15 min of harvest using 4% paraformaldehyde for 15 min, and permeabilized with 0.2% Triton X-100 for 10 min. After blocking in 5% normal serum, cells were incubated overnight at 4°C with a validated monoclonal antibody against 8-nitroguanine, followed by an Alexa Fluor-conjugated secondary antibody. Fluorescence was quantified by flow cytometry and expressed as background-subtracted median fluorescence intensity and fold change over untreated cells.

### DNA damage response and apoptosis

Phosphorylated H2AX was detected using rabbit monoclonal anti-phospho-Histone H2A.X Ser139 antibody (Cell Signaling Technology, 9718, clone 20E3, RRID: AB_2118009). Cells were fixed in 4% paraformaldehyde for 15 min, permeabilized using 90% ice-cold methanol for 30 min, incubated with primary antibody at 1:200 for 1 h at room temperature, and detected using an Alexa Fluor 647-conjugated anti-rabbit secondary antibody. γH2AX was expressed as both the percentage of γH2AX-positive viable singlets and median fluorescence intensity. Apoptosis was measured using the APC Annexin V Apoptosis Detection Kit with 7-AAD (BioLegend, 640930). Unfixed cells were stained in annexin-binding buffer for 15 min at room temperature protected from light and analyzed within 1 h. Early apoptotic cells were defined as annexin V-positive/7-AAD-negative, and late apoptotic or membrane-compromised cells as annexin V-positive/7-AAD-positive. Caspase-3/7 activity was measured in parallel cultures using the Caspase-Glo 3/7 Assay (Promega, G8091). An equal volume of reagent was added to cells, plates were mixed for 30 s and incubated for 30 min at room temperature, and luminescence was measured. Caspase activity was normalized to viable-cell number in replicate wells and expressed as fold change relative to untreated cells. All flow-cytometric measurements were performed on a MACSQuant Analyzer 10 (Miltenyi Biotec; RRID: SCR_020268). Compensation was established using single-color controls, gates were defined using fluorescence-minus-one or secondary-only controls where appropriate, and at least 10,000 viable singlet events were acquired per sample.

### Metabolomics

Liquid Chromatography Mass Spectrometry (LC-MS) was performed for metabolomics using a Vanquish Ultra High-Performance LC system (ThermoFisher Scientific, RRID: SCR_020943) coupled to an Orbitrap ID-X Tribrid mass spectrometer (ThermoFisher Scientific, RRID: SCR_021134), equipped with a heated electrospray ionization (H-ESI) source capable of both positive and negative modes simultaneously. Before analysis, the MS instrument was calibrated using the Pierce FlexMix Calibration Solution (ThermoFisher Scientific, Cat# A39239, RRID: AB_2801668). Cell or tissue samples, along with blank controls, were placed in the autosampler. Individual samples were run alongside a pooled sample made from an equal mixture of all individual samples to ensure chromatographic consistency. Chromatographic separation of metabolites was achieved by hydrophilic interaction liquid chromatography (HILIC) using a Luna® 3 µm NH₂ column, 150 mm × 2.1 mm (Phenomenex, Cat# 00F-4378-B0, RRID: AB_2890953) in conjunction with a SecurityGuard™ HILIC guard column (Phenomenex, Cat# AJ0-4287, RRID: AB_2890954). The column temperature and flow rate were maintained at 27 °C and 0.2 mL/min, respectively. Mobile phases consisted of A (5 mM ammonium acetate, 48.5 mM ammonium hydroxide, pH 9.9) and B (100% acetonitrile). The following linear gradient was applied: 0.0-0.1 min: 85-80% B, 0.1-17.0 min: 80-5% B, 17.0-24.0 min: 5% B, 24.0-25.0 min: 5-85% B, 25.0-36.0 min: 85% B. The injection volume and autosampler temperature were kept at 5 µL and 4 °C, respectively. High-resolution MS was acquired using a full scan method alternating between positive and negative polarities (spray voltages: +3800 kV/-3100 kV; sheath gas flow: 45 arbitrary units; auxiliary gas flow: 15 arbitrary units; sweep gas flow: 1 arbitrary unit; ion transfer tube temperature: 275 °C; vaporizer temperature: 300 °C). Three mass scan events were set for the duration of the 36-minute run time: (1) negative polarity full scan, 70-975 m/z; (2) positive polarity full scan, 70-360 m/z; (3) positive polarity full scan, 360-1500 m/z. MS1 data were acquired at a resolution of 60,000 with a standard automatic gain control and a maximum injection time of 100 ms.

Data was acquired using Xcalibur software (ThermoFisher Scientific, RRID: SCR_014593). Chromatograms were reviewed using FreeStyle software (ThermoFisher Scientific, RRID: SCR_016664) with a 5-ppm mass tolerance. Peak areas were quantified using TraceFinder software (ThermoFisher Scientific, RRID: SCR_016602) from either positive or negative modes, depending on previously run standards. Peak areas were corrected to blank samples and normalized to protein content and the total ion count of that sample.

### In vivo studies

All studies were approved by the University of California, San Francisco Institutional Animal Care and Use Committee (IACUC). Cells were intracranially injected into the cortex of 5-6 weeks old female C57BL/6 mice (SB28mut) or CB17 SCID mice (133-A6, 130-A7, TS603, SF10417, BT142)^26^. Intracranial implantation was performed using a stereotactic frame at 2 mm to the right of the medial suture, 2 mm behind the bregma, and a depth of 2 mm. Tumor volume was determined by BLI (SB28, 133-A6, 130-A7) or T2-weighted MRI using a Bruker 3T scanner (TS603). Tumor growth was monitored by BLI on a Xenogen IVIS Spectrum imaging system (PerkinElmer/Caliper Life Sciences, RRID: SCR_018520) 10 min after intraperitoneal injection of D-Luciferin, Potassium Salt (Gold Biotechnology, Cat# LUCK-100, RRID: SCR_018804) prepared according to the manufacturer’s directions. The total flux signal was used for quantification in all studies. Axial T2-weighted images were acquired using a spin-echo TurboRARE sequence (TE/TR = 64/3700 ms, FOV = 30 x 30 mm^2^, 256 x 256, slice thickness = 1.5 mm, NA = 4). Once tumors were detectable, mice bearing SB28mut or 133-A6 tumors were treated with vehicle (Oraplus), VOR (50 mg/kg/day, 6 days/week in Oraplus) by oral gavage, LIR (0.4 mg/kg/day, 6 days/week in PBS) by intranasal instillation, or the combination of VOR and LIR as above. Mice bearing 130-A7 tumors were treated with vehicle (Oraplus), VOR (50 mg/kg/day, 6 days/week in Oraplus) by oral gavage, NLY01 (0.4 mg/kg/day, 6 days/week in PBS) by intranasal instillation, or the combination of VOR and NLY01. Mice were treated and monitored for survival until they needed to be euthanized according to IACUC guidelines.

## ACKNOWLEDGEMENTS

The authors acknowledge support from the Preclinical NMR Core of the Department of Radiology and Biomedical Imaging at UCSF. The schematics in this paper were created with BioRender. We thank Neuraly Inc. for providing NLY01 for our studies. We thank Dr. Timothy Phoenix (University of Cincinnati), Dr. Timothy Chan (Cleveland Clinic), Dr. Joseph Costello (University of California, San Francisco) and Dr. Hideho Okada (University of California, San Francisco) for providing the mIDH glioma models used in this study.

## FUNDING

These studies were funded by the following grants: NIH R01CA288389 (PV), the American Cancer Society Research Scholar Grant (PV; RSG-24-1258893-01-CDP), the UCSF Loglio Initiative (PV, JDG), and the Gianna Rae Meadows fund for the Oligodendroglioma Cure (PV, JDL). We acknowledge support from the NIH S10OD030256 grant.

## AUTHOR CONTRIBUTIONS

PV conceived the project. PV, CT and SU developed the methods, performed experiments, and analyzed the data. PV wrote the original manuscript. TP provided cell lines, JDL and JDG provided intellectual input. PV supervised the project and was responsible for management and coordination. PV, JDL, and JDG acquired funding.

## COMPETING INTERESTS

The authors declare that they have no competing interests.

## DATA AND MATERIAL AVAILABILITY

All data needed to assess this manuscript are available in the main paper or the Supplementary Information.

## Notes

### Competing Interest Statement

The authors have declared no competing interest.

