## Supplementary Figures S1-S6 for "GLP1R agonists and vorasidenib converge on peroxynitrite-mediated cytotoxicity in mutant IDH glioma"

Supplementary Figure S1

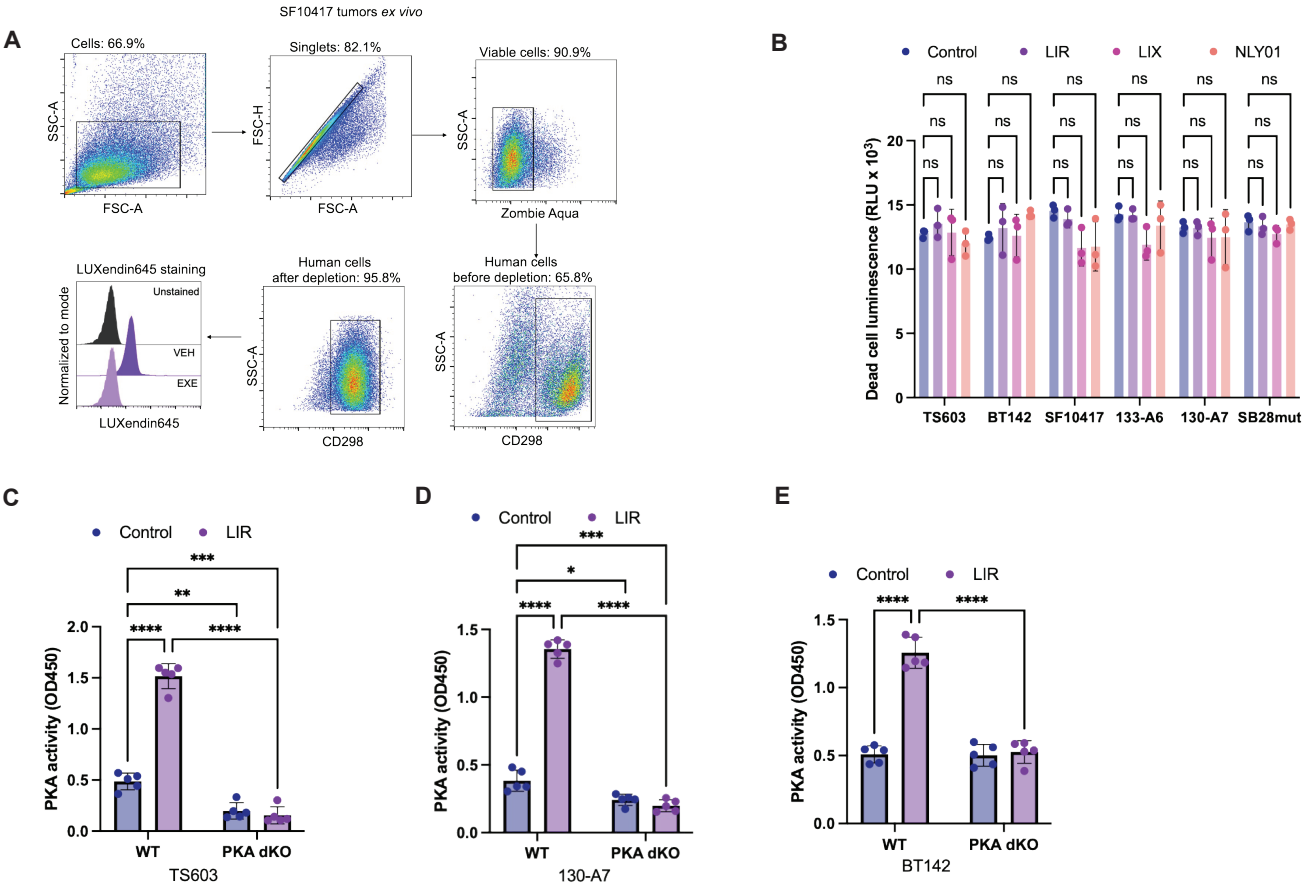

### SUPPLEMENTARY FIGURE LEGENDS

**Supplementary Figure S1. GLP1R labeling in resected tumors, absence of GLP1R agonist-induced cell death, and PKA dependence of the anti-proliferative effect. (A)** Representative gating for analysis of tumors resected from mice bearing intracranial SF10417 tumors: forward and side scatter to identify cells, FSC-H against FSC-A to exclude doublets, Zombie Aqua to exclude non-viable cells, and CD298 to identify human tumor cells before and after depletion of mouse stromal cells, with the percentage of events retained at each step indicated. The histogram shows LUXendin645 labeling of CD298-positive cells after vehicle (VEH) or exendin-4(9-39) (EXE) competition, with unstained cells as the background reference; quantification of this experiment is shown in Fig. 1D. **(B)** Cell death, measured as dead-cell luminescence (RLU $\times 10^3$), in TS603, BT142, SF10417, 133-A6, 130-A7 and SB28mut cells treated with vehicle (Control), LIR, LIX or NLY01 for 72 h. None of the agonists increased cell death in any model. **(C-E)** PKA activity (OD450) in parental (WT) and PKA double-knockout (PKA dKO) **(C)** TS603, **(D)** 130-A7 and **(E)** BT142 cells treated with vehicle (Control) or LIR for 72 h. Data are mean ± SD with individual biological replicates shown ($n \geq 3$ per group). \*\*\*\*$p < 0.0001$, \*\*\*$p < 0.001$, \*\*$p < 0.01$, \*$p < 0.05$; ns, not significant.

**Supplementary Figure S2**

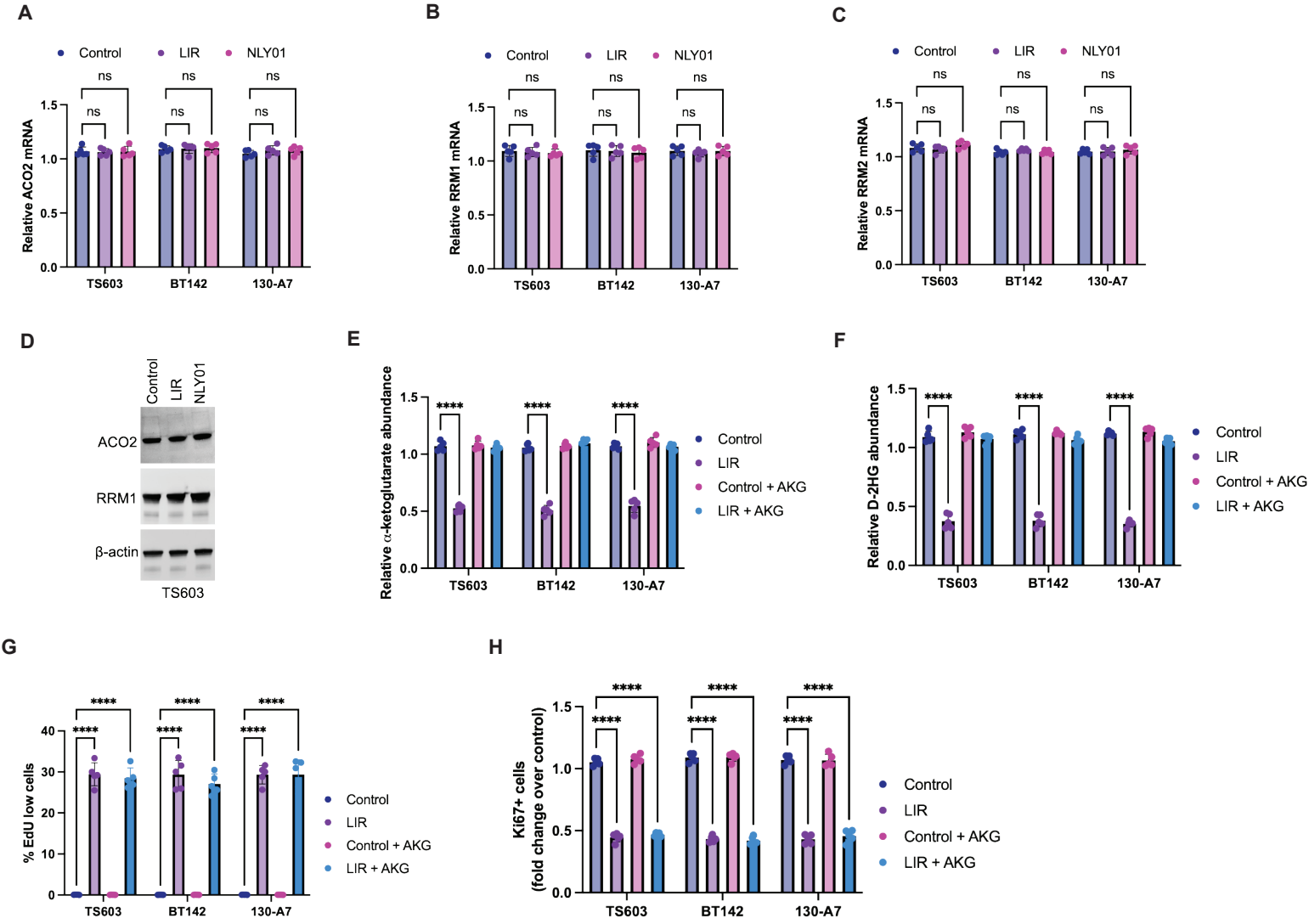

**Supplementary Figure S2. GLP1R agonists do not alter ACO2, RRM1, or RRM2 abundance. (A-C)** Relative ACO2 **(A)**, RRM1 **(B)** and RRM2 **(C)** mRNA in TS603, BT142 and 130-A7 treated with vehicle, LIR or NLY01. **(D)** Immunoblot for ACO2, RRM1, and beta-actin in TS603 cells treated with vehicle, LIR, or NLY01. **(E-H)** Relative α-ketoglutarate **(E)**, D-2HG **(F)**, EdU-low cells as a percentage of total **(G)**, and Ki67+ cells (fold change over control; **H**) in TS603, BT142, and 130-A7 treated with vehicle or LIR, with or without cell-permeable α-ketoglutarate (AKG). Data are mean ± SD with individual biological replicates shown ($n \geq 3$ per group). \*\*\*\*$p < 0.0001$, \*\*\*$p < 0.001$, \*\*$p < 0.01$, \*$p < 0.05$; ns, not significant.

Supplementary Figure S3

A

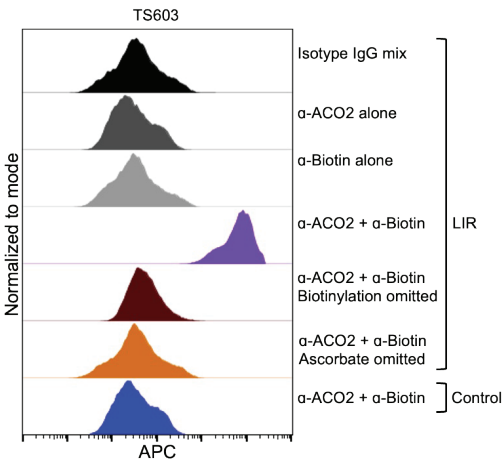

B

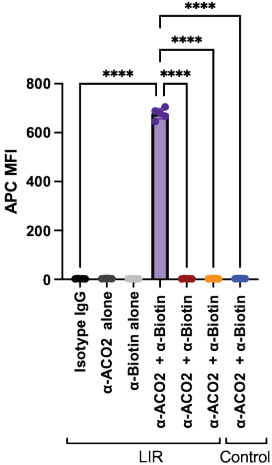

C

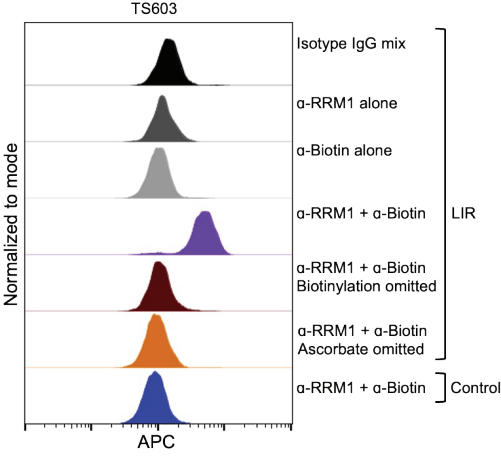

D

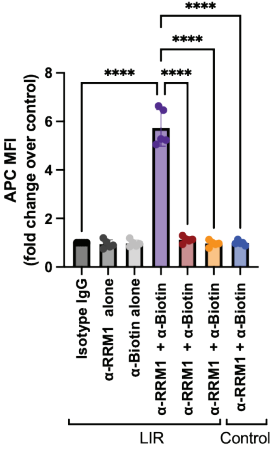

E

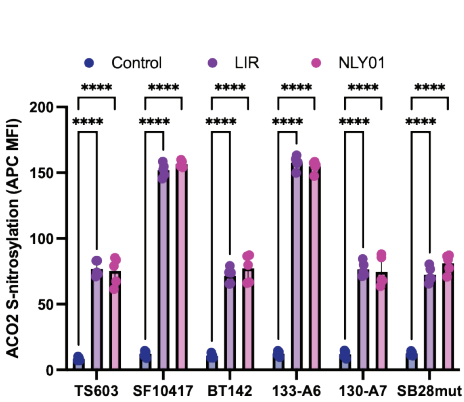

F

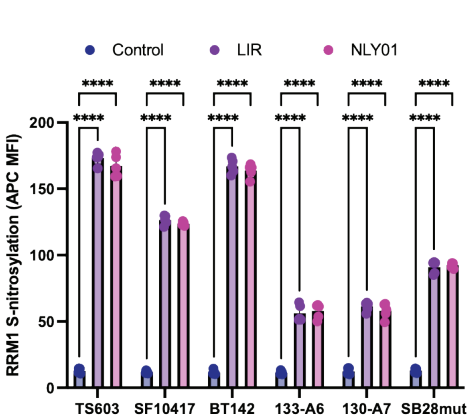

G

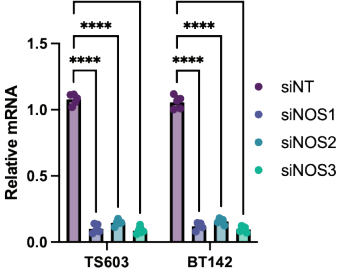

H

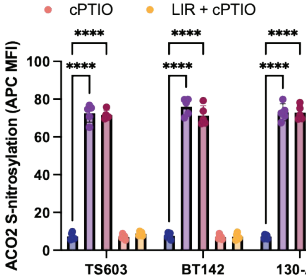

I

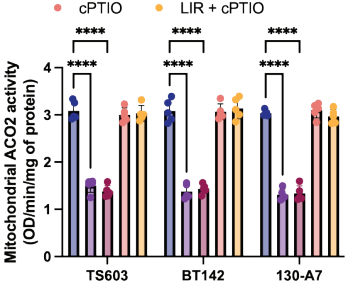

**Supplementary Figure S3. Specificity controls for the biotin-switch proximity ligation assay, and extended S-nitrosylation and NOS data.** (A) Representative biotin-switch PLA signal for ACO2 in LIR-treated TS603: isotype-matched IgG for both primary antibodies; anti-ACO2 alone; anti-biotin alone; the complete reaction; the complete reaction with biotinylation omitted; the complete reaction with ascorbate omitted; and the complete reaction in untreated cells. All arms except the last are in the LIR background. (B) Quantification of A as APC MFI. (C) The equivalent control set for RRM1 in LIR-treated TS603. (D) Quantification of C as APC MFI, expressed as fold change over the untreated control. (E, F) ACO2 (E) and RRM1 (F) S-nitrosylation by biotin-switch PLA (APC MFI) in six mIDH glioma models treated with vehicle, LIR or NLY01. (G) Relative mRNA in TS603 and BT142 transfected with siNT, siNOS1, siNOS2 or siNOS3; each bar reports the transcript targeted by the corresponding siRNA. (H) ACO2 S-nitrosylation (APC MFI) in TS603, BT142 and 130-A7 treated with vehicle, LIR, NOC-18, cPTIO, or LIR with cPTIO. (I) Mitochondrial ACO2 activity in the same cells and arms. Panel B reports raw APC MFI and panel D fold change over the untreated control, so the two y-axes are not comparable; the panels are also from separate acquisitions and are not intended for direct comparison. Flow cytometry histograms are normalized to mode and plotted on an identical axis range. Data are mean ± SD with individual biological replicates shown ($n \geq 3$ per group). \*\*\*\*$p < 0.0001$, \*\*\*$p < 0.001$, \*\*$p < 0.01$, \*$p < 0.05$; ns, not significant.

Supplementary Figure S4

A

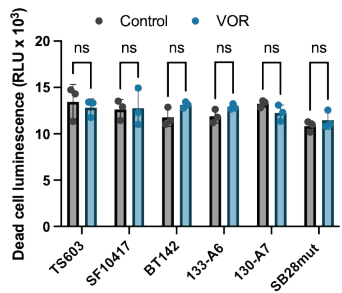

B

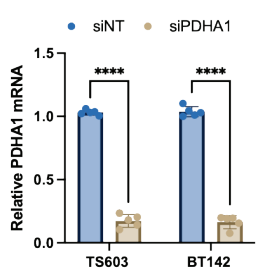

**Supplementary Figure S4. VOR is not cytotoxic as a single agent, and PDHA1 knockdown validation. (A)** Dead-cell luminescence (RLU $\times 10^3$) in six mIDH glioma models treated with vehicle or VOR. **(B)** Relative PDHA1 mRNA in TS603 and BT142 transfected with siNT or siPDHA1. Data are mean ± SD with individual biological replicates shown ($n \geq 3$ per group). \*\*\*\*$p < 0.0001$, \*\*\*$p < 0.001$, \*\*$p < 0.01$, \*$p < 0.05$; ns, not significant.

Supplementary Figure S5

A

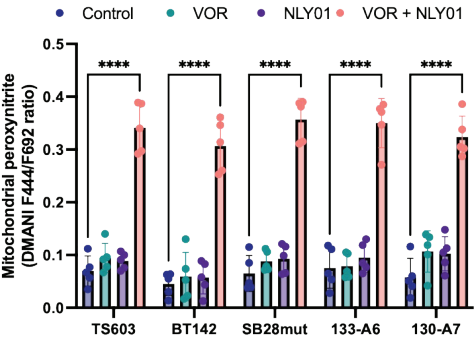

B

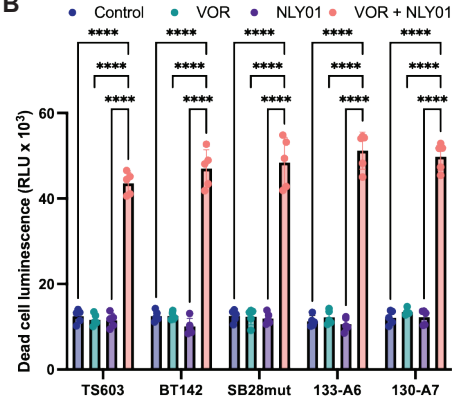

**Supplementary Figure S5. The NLY01 combination reproduces the peroxynitrite and cytotoxicity phenotype.** **(A)** Mitochondrial peroxynitrite (DMANI F444/F692 ratio) in TS603, BT142, SB28mut, 133-A6 and 130-A7 treated with vehicle, VOR, NLY01, or VOR with NLY01. **(B)** Dead-cell luminescence (RLU x 10<sup>3</sup>) in the same models and arms. Data are mean ± SD with individual biological replicates shown (n ≥ 3 per group). \*\*\*\*p<0.0001, \*\*\*p<0.001, \*\*p<0.01, \*p<0.05; ns, not significant.

Supplementary Figure S6

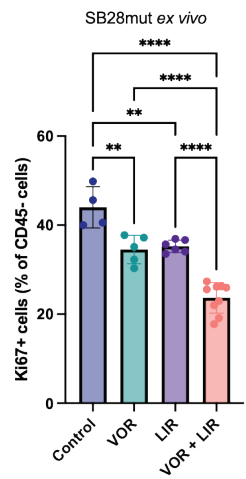

**Supplementary Figure S6. Tumor cell proliferation in SB28mut tumors.** Ki67-positive cells as a percentage of CD45-negative cells in SB28mut tumor tissue resected at endpoint from mice treated with vehicle (Control), VOR, LIR, or the combination of VOR and LIR. Representative histograms are shown in Fig. 7D. Data are mean ± SD with individual biological replicates shown ($n \geq 3$ per group). \*\*\*\*$p < 0.0001$, \*\*\*$p < 0.001$, \*\*$p < 0.01$, \*$p < 0.05$; ns, not significant. Gating on CD45-negative cells excludes tumor-infiltrating leukocytes, so the readout reports proliferation of tumor cells rather than of the whole dissociated tumor.
